# Hfq Globally Binds and Destabilizes the bound sRNAs and mRNAs in *Yersinia pestis*

**DOI:** 10.1101/487769

**Authors:** Yanping Han, Dong Chen, Yanfeng Yan, Hongduo Wang, Zizhong Liu, Yaqiang Xue, Ruifu Yang, Yi Zhang

## Abstract

Hfq is a ubiquitous Sm-like RNA binding protein in bacteria involved in physiological fitness and pathogenesis, while its *in vivo* binding natures still remain elusive. Here we reported the first study of the Hfq-bound RNAs map in *Yersinia pestis*, the causative agent of a kind of plague, by using Cross-Linking Immunoprecipitation coupled with deep sequencing (CLIP-Seq) approach. We show that Hfq binds over 80% mRNAs of *Y. pestis*, and also globally binds non-coding sRNAs encoded by the intergenic, antisense, and the 3’ regions of mRNAs. Hfq U-rich stretch is highly enriched in sRNAs, while motifs partially complementary to AGAAUAA and GGGGAUUA are enriched in both mRNAs and sRNAs. Hfq binding motifs are enriched at both terminal sites and in the gene body of mRNAs. Surprisingly, a large fraction of the sRNA and mRNA regions bound by Hfq and those downstream are destabilized, likely via a 5’P-activated RNase E degradation pathway and consistent with Hfq-facilitated sRNA-mRNA base-pairing and the coupled degradation in *Y. pestis*. These results together have presented a high-quality Hfq-RNA interaction map in *Y. pestis*, which should be important for further deciphering the regulatory role of Hfq-sRNAs in *Y. pestis*.

**AUTHOR SUMMARY:** Discovered in 1968 as an *Escherichia coli* host factor that was essential for replication of the bacteriophage Qβ, the Hfq protein is a ubiquitous and highly abundant RNA binding protein in many bacteria. Under the assistance of Hfq, small RNAs in bacteria play important role in regulating the stability and translation of mRNAs by base-pairing. In this study, we want to elucidate the Hfq assisted sRNA-mRNA regulation in *Yersinia pestis*. A global map of Hfq interaction sites in *Y. pestis* was obtained by sequencing of cDNAs converted from the Hfq-bound RNA fragments using UV cross-linking coupled immunoprecipitation technology. We demonstrate that Hfq could hundreds of sRNAs and the majority of mRNAs in living *Y. pestis*. The enriched binding motifs in sRNAs and mRNA are significantly complementary to each other, suggesting a general base-pairing mechanism for sRNA-mRNA interaction. The Hfq-bound sRNA and mRNA regions were both destabilized. The results suggest that Hfq binding facilitates sRNA-mRNA base-pairing and coordinates their degradation, which might enable Hfq to surveil the hemostasis of most mRNAs in bacteria.

## INTRODUCTION

Hfq is a small, ubiquitous and highly abundant protein in bacteria involved in physiological fitness and pathogenesis [1]. It forms a doughnut-like homohexameric structure and belongs to Sm/LSm superfamily of RNA-binding proteins [2, 3]. Hfq is the only bacterial LSm homolog. Similar to its eukaryotic counterparts, each monomer carries the signature Sm motif for protein-protein interaction and RNA binding that contribute to post-transcriptional regulation [2, 4, 5].

The largest class of regulatory RNAs in bacteria is small RNA transcripts including the cis-encoded antisense sRNAs and trans-encoded sRNAs. Antisense sRNAs have extensive potentials for base-pairing with their target RNA, while the trans-encoded sRNAs have more limited complementarity. Both subclasses of sRNAs are capable of causing translation inhibition, mRNA cleavage or degradation.

Hfq plays a central role in mediating sRNA regulation of mRNA stability in bacteria [6, 7]. Hfq binds to sRNAs and promotes the limited base-pairing with their mRNA targets [2–10]. Hfq-promoted sRNA binding target sites are not only located in the canonical Shine-Dalgarno SD/AUG region, but also in other regions of the target mRNAs [11–13].

A key endoribonuclease for RNA processing and decay in gamma *proteobacteria* is RNase E, which recognizes its substrates via two different modes of action [14]. RNase E senses and binds the 5’ monophosphate group of a target, which enables the enzyme to distinguish and prefer broken RNAs that have already undergone at least one ribonuclease cleavage from the primary transcript with a 5’ triphosphate group. The cleavage sites activated by the 5’-monophosphate binding can be quite downstream [15]. Alternatively, RNase E directly makes a cut in the body of an mRNA [4, 5, 14]. RNase E is crucial for sRNA-induced decay of target mRNAs and sRNAs themselves [4, 14–16]. RNase E degradation of sRNAs can occur either when they are free, or when paired with their targets. The latter is called the coupled degradation [16–18]. RNase E contains a defined region for Hfq binding [19]. The interaction between RNase E and Hfq seems to play an important role in sRNA-mediated mRNA decay [4, 14]. However, it is yet unclear how much does this potential novel type of “degradosome” acts in controlling bacterial mRNA decay.

Hfq is known to bind most sRNAs via its proximal face binding of the polyU sequences typical of Rho-independent terminators, with uridine stacked in pockets between neighboring monomers around the central pole [10, 20–24]. The distal face binds various A-rich sequences located in mRNAs and sRNAs Hfq-mRNA interactions [25–27]. It has recently been proved that the rim (lateral face) of Hfq contacts UA-rich sequences in sRNA and mRNA [10, 28–32]. Hfq can simultaneously bind sRNA and mRNA to form Hfq-RNA ternary complexes [22, 29], probably via the engagement of multiple Hfq surfaces. The multiple binding surfaces of an Hfq homohexamer enable a large flexibility of this protein in mediating not only the sRNA-mRNA interaction, but also probably the sRNA-sRNA interactions in regulating the stabilities of both types of bacterial RNAs [10, 24].

Genome-wide analysis of Hfq-bound *in vivo* RNA fragments in *Escherichia coli* has revealed the enrichment of A-R-N (A-A/G-any nucleotide) associated with the Shine-Dalgarno translation initiation sequence in mRNAs [33], suggesting a prevalence of Hfq binding of bacterial mRNAs.

The mechanism of Hfq-mediated RNA regulation has been extensively studied in model gram-negative bacteria *E. coli,* but not well-studied in gram-negative pathogenic bacteria. Plague caused by *Y. pestis* is a zoonotic disease primarily transmitted between fleas and mammals. Hfq is found to be required for the virulence of *Y. pestis* and other *Yersinia* species [34, 35]. The *Y. pestis hfq* gene (also annotated as *ymr*) encodes an 101 amino acid Hfq homolog, which shared ∼85% similarity with their homologues in *E. coli* and *S. typhimurium*, showing 100% sequence identity within the Sm1 and Sm2 signature regions [2].

In this study, we sequenced and analyzed cDNA libraries generated from the Hfq-bound *Yersinia pestis* RNA fragments using two different UV cross-linking and immunoprecipitation methods, one resembling CLIP-seq and another resembling RIP-seq. Considering that Hfq normally binds single stranded region near a hairpin structure, we mildly digested Hfq-bound RNA molecules in this study to recover RNA segments containing Hfq-bound sites. We have analyzed cDNA reads from Hfq-bound RNA after anti-FLAG cross-linking immunoprecipitation in *E. coli* [33]. However, their reads were too short to give sufficient peak information, let alone the motif information (data not shown). Such a problem was effectively eliminated from this study. Using transcriptome sequencing data as controls, we showed that Hfq binds most of the expressed mRNAs and sRNAs in bacterial cells. Hfq binds mRNAs not only via known motifs, but also via the previously unknown G-rich motifs. Moreover, we demonstrated an increased destabilization of RNA segments that are bound by Hfq, irrespective of whether they are located in sRNAs or in mRNAs. The results suggest that the Hfq-facilitated sRNA-mRNA base pairing might be more likely coupled with their degradation than previously appreciated.

## RESULTS

### CLIP-Seq Revealed that Hfq Binds over 80% of the Transcribed mRNAs in *Y. pestis*

To globally map Hfq binding RNA and binding sites in *Y. pestis* living cells, we used two strains expressing a FLAG epitope. The experimental Hfq strain expressed Hfq-FLAG from a plasmid in ΔHfq genetic background. The control strain was a wild-type *Y. pestis* strain expressing truncated Hfq fused with FLAG epitope. The growth study showed that the exogenously expressed Hfq-FLAG was functionally competent (data not shown). Using native polyacrylamide gel electrophoresis (PAGE), we found that Hfq-FLAG stably existed as trimer and hexamer in bacterial cells (Fig S1A). RNA-Seq profiling of the transcriptomes of the constructed strains demonstrated a very similar genome expression pattern of these two strains (Fig S1B).

The FLAG tagged Hfq-RNA complex were cross-linked by UV irradiation of cultured cells, followed by co-immunoprecipitation by anti-FLAG and partial digestion of unprotected RNA segments by RNase T1 (Fig S2 for experimental strategy). The Hfq-bound RNA segments were purified and ligated with adaptors for Illumina sequencing (Fig S1C). Equal amount of bacterial cultures from the Hfq-FLAG and WT-Flag strains were lysed and subjected to parallel immunoprecipitation experiments to obtain our CLIP-Seq data. Experiments on these two strains were strictly performed in parallel. However, we generally obtained much less cDNAs from WT-FLAG strain compared to the Hfq-FLAG strain, suggesting that FLAG tag itself did not yield much RNA binding noise and therefore was a successful co-IP system (Fig S1C). We obtained 18.1 million of Hfq-FLAG-bound RNA tags and 2.1 million of FLAG-bound control tags mapped onto the *Y. pestis* 91001 genome (Table S1, Fig 1C). Free FLAG control exclusively bound rRNA (86.03%) and tRNA (2.38%), indicative of the non-specific binding (Table S2). The fraction of CLIP-Seq reads from the Hfq-FLAG sample mapped to the annotated sRNA region was 10-fold of the FLAG control, consistent with the specific sRNA binding activity of Hfq. Meanwhile, Hfq-FLAG sample also demonstrated 8.17-fold and 14.23-fold increased binding in the mRNA and intergenic regions respectively (Table S2). The increased Hfq binding in sRNA, mRNA and intergenic regions seems to be genome-wide rather than to some specific genes (Fig 1A). These results suggest that Hfq selectively binds a large population of sRNA and mRNA in living bacterial cells. The enriched binding of RNA transcribed from intergenic region is consistent with the assumption that intergenic regions harbor a large number of un-annotated sRNAs, which is further described below.

**Fig 1.**
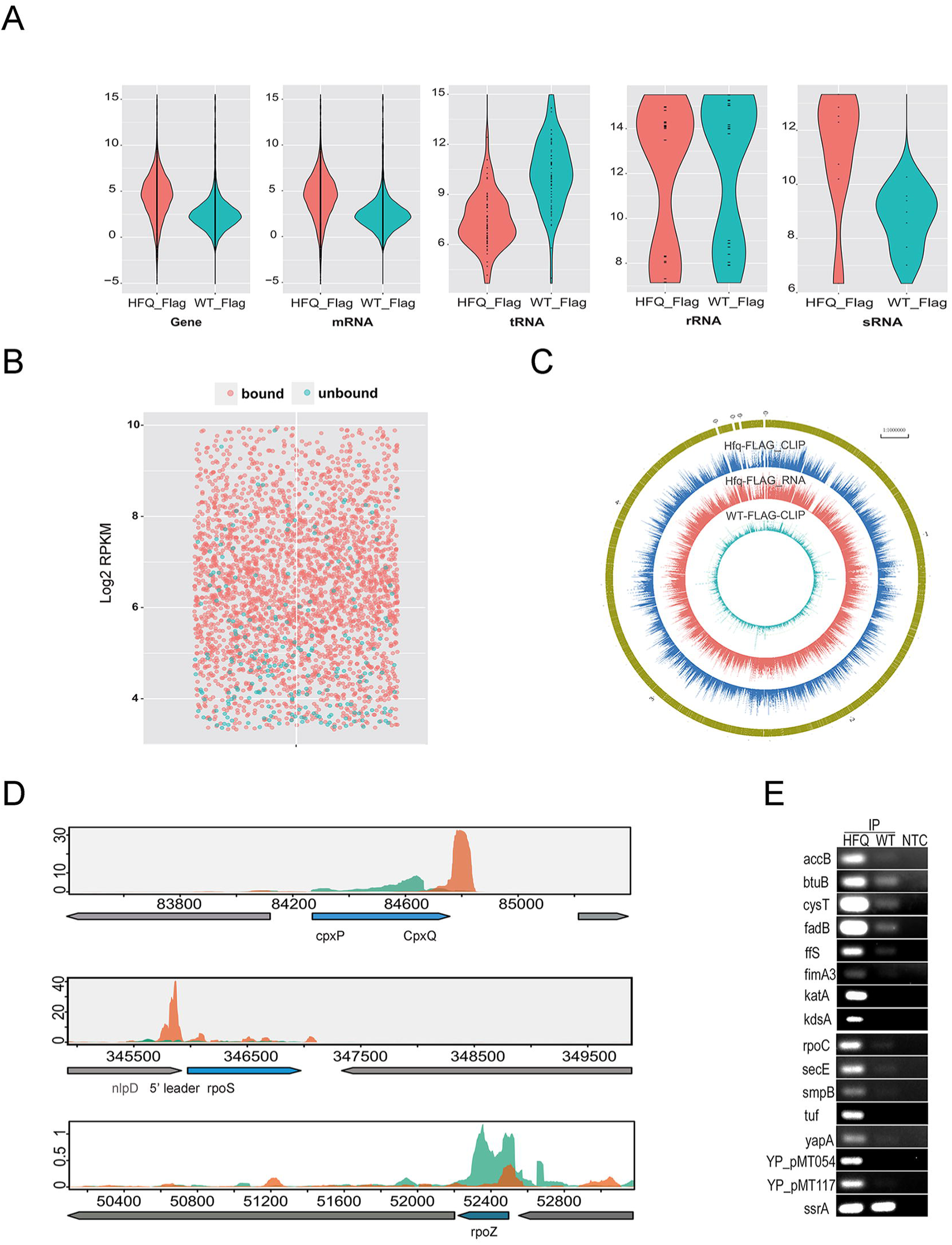
CLIP-Seq revealed that Hfq binds over 80% of transcribed mRNAs in *Y. pestis.* (A) Violin plot of Hfq binding profiles in all genes, mRNA, tRNA, rRNA and seven annotated sRNAs. Red violin represents *Hfq* Flag strain, and blue for WT Flag strain. (B) Dot plot of gene expression level classified by Hfq binding. Red dots represent bound genes and blue for unbound genes. (C) Reads density presentation of whole genome for three sequencing sample, RNA-Seq of Hfq Flag, CLIP-Seq of Hfq Flag, and CLIP-Seq of WT Flag, from external to internal respectively. (D) Reads density plot for Hfq-bound *cpxP* and *rpoS* mRNA, and for Hfq-unbound *rpoZ* in *Y. pestis*. Orange represents Hfq binding density, and seagreen for RNA expression density. (E) RIP-PCR validation of Hfq-bound sRNA and mRNAs.

To identify the Hfq-bound genes from our CLIP-Seq data, we normalized the CLIP-Seq reads in each gene to the non-specific bound 23S rRNA gene *YP_r2*. This rRNA represents the most abundant one among all CLIP identified RNAs in both strains. The analysis reveals a total of 3331 Hfq-bound and 864 Hfq-unbound genes in *Y. pestis* (Table S3). All of the 22 rRNA genes were Hfq-unbound, and 65 out of 68 tRNA genes were Hfq-unbound. Among the 7 annotated sRNAs, 6 were detected by the CLIP-Seq reads, 4 were identified as Hfq-bound and 2 were Hfq-unbound (Table S4). The Hfq-bound sRNAs includes the well-studied Spf, Ffs, CsrB and ssrS. In previous studies [36, 37], Spf sRNA is Hfq-bound in both *E. coli* and *Samonella*. Ffs and SsrB are not bound by Hfq in *E coli*, and csrB and ssrS are not Hfq-bound in *Samonella*. The Hfq-unbound sRNAs in *Y. pestis* included SsrA and RnpB. Both are Hfq-unbound in *E. coli* (Table S4).

We showed that 80.5% (3323 out of 4128) of all mRNA genes were Hfq-bound, which is at a higher frequency than sRNAs. Transcriptome sequencing data from the two experimental *Y. pestis* strains cultured under the same condition were obtained as another set of controls (Table S1). The CLIP method has detected that Hfq-bound and unbound genes were well expressed (Fig 1B). Interestingly, it seems that Hfq-bound genes tend to be clustered in the lower expressed gene population (Fig 1B). Hfq-bound genes were clustered in a large array of metabolic pathways related to the *in vitro* exponential growth, while Hfq-unbound genes were enriched in flagellar assembly, bacterial secretion system and chemotaxis which are important host infection. These results collectively suggested that Hfq binds to most genes important for the exponential growth of *Y. pestis*, which supports its global and extensive regulatory role (Fig S1D).

Comparison between Hfq binding profiles in the *Y. pestis* genome and the corresponding RNA-Seq reads indicated the binding specificity. For example, the CLIP-Seq and RNA-Seq reads peaked at different locations for the previously known Hfq-bound *cpxP* mRNA. The RNA-Seq reads peaked at the coding region, while CLIP-Seq reads peaked at the 3’UTR region corresponding to the CpxQ sRNA (Fig 1D, upper). The Hfq binding of CpxQ sRNA has been recently reported to play a role in protecting bacteria against inner membrane damage [38]. The 5’ leader of *rpoS* mRNA is located at the 3’ part of *nlpD* mRNA, and is known to be bound by Hfq and DsrA sRNA [22, 39]. We showed that Hfq has a strong binding peak in the 5’ leader region of rpoS mRNA. Moderate binding peaks in the gene body region and 3’ downstream region were also evident (Fig 1D, middle). The CLIP-Seq and RNA-Seq profiles of one example of Hfq-unbound mRNAs, *rpoZ* were also shown (Fig 1D, bottom).

We validated the Hfq-bound mRNA and sRNAs obtained above using RIP-PCR analysis. After co-immunoprecipitation by anti-FLAG using Hfq-FLAG and WT-FLAG cell lystate, we used SsrA sRNA as an unbound control to validate 15 randomly chosen Hfq-bound sRNAs and mRNAs, including Ffs, *btuB*, *cysT*, *fadB*, *fimA3*, *katA*, *kdsA*, *secE*, *smpB*, *tuf*, *yapA*, *YP-PMT054* and *YP-PMT117*. All of these Hfq-bound and unbound sRNAs and mRNAs were validated (Fig 1E).

### RIP-Seq Experiments Confirmed the Ubiquitous Nature of Hfq Binding of mRNAs

To more globally validate the Hfq CLIP-Seq results, we performed two independent sets of Hfq RIP-Seq experiments with the same *Y. pestis* strains and experimental conditions. A total of 9,186,327 and 26,971,805 clean reads were obtained from the Hfq-FLAG strains. The mapping features of RIP-Seq reads were strikingly similar to that of the CLIP-Seq data (Table S1). Moreover, distribution of RIP-Seq reads in all genes was more similar between the two repeated experiments and with the CLIP-Seq reads when compared to their correlation with RNA-Seq and FLAG controls (Fig 2A, Fig S3A). When Hfq-bound and unbound genes were similarly identified from the two sets of RIP-Seq data, the results showed that the Hfq-bound mRNAs and sRNAs were highly overlapped among different immunoprecipitation experiments (Fig 2B).

**Fig 2.**
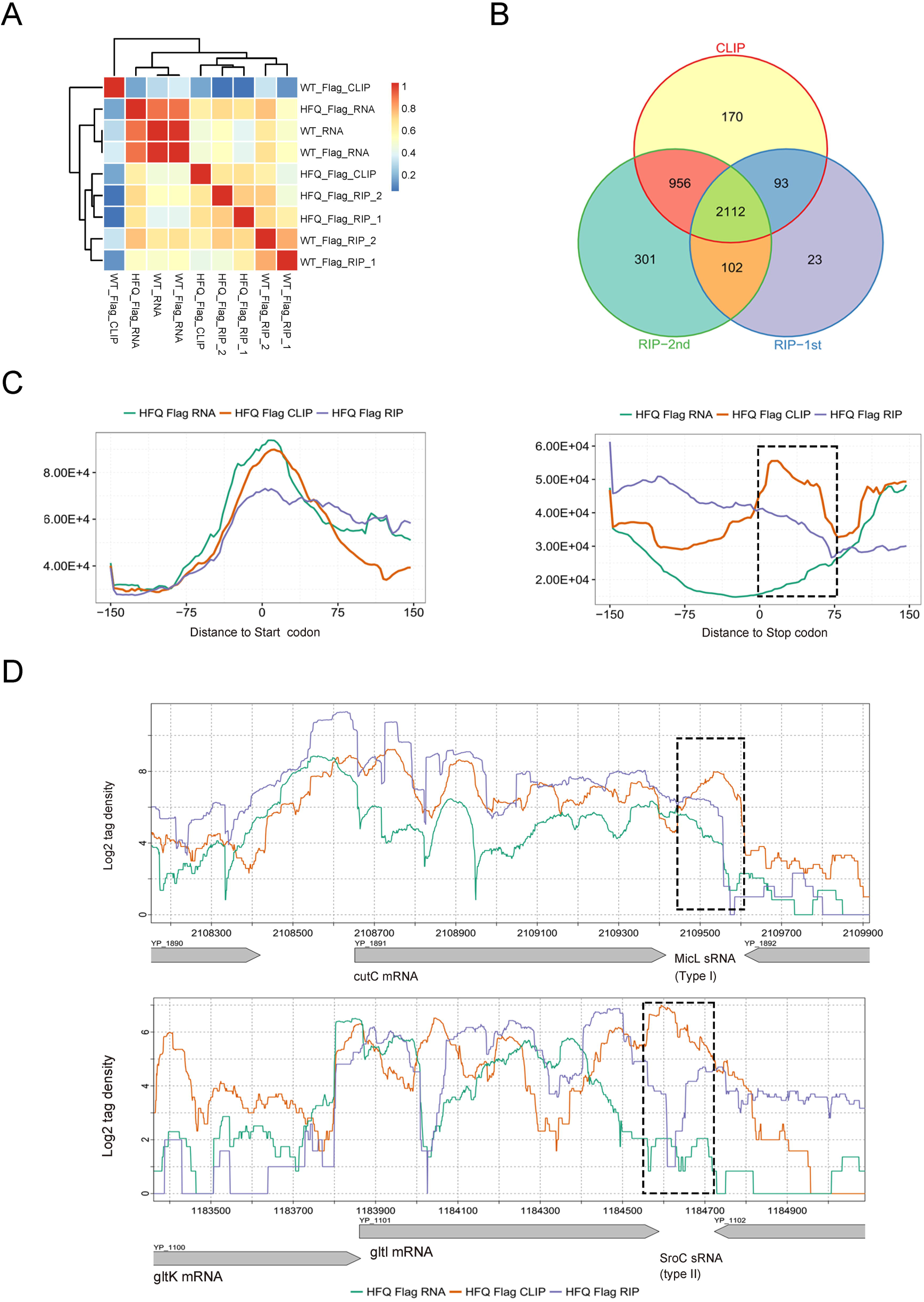
RIP-Seq experiments confirmation of the global binding properties of Hfq. (A) Pearson correlation analysis of RNA-Seq (one set), CLIP-Seq (one set) and RIP-Seq (two sets) experiments using different strains. (B) Venn diagram of the Hfq-bound mRNAs and sRNAs among three different immunoprecipitation experiments. (C) Reads distribution around the 5’ leader region and 3’ region from the CLIP-Seq and RNA-Seq data and one set of RIP-Seq data. Left panel represents for 5’ leader regions and right panel for 3’ regions. Black dashed box indicates the region downstream of stop codon and enriched in CLIP peak. (D) Reads density illustration of two major sRNA types transcribed from mRNA 3’ region. Top panel represents a type I sRNA and bottom panel for a type II sRNA. Black dashed box represents the location of predicted sRNAs.

### CLIP-seq and RIP-Seq data Suggest a Global Regulatory Role Encoded by Intergenic and Antisense RNAs, and also by the 3’ Regions of mRNAs

In order to better understand the length features of Hfq-bound intergenic and antisense RNAs revealed by CLIP-Seq data, longer Hfq-bound RNA segments (insert showing a gel mobility longer than 150nt) were selected for sequencing in RIP-Seq experiments shown in this study. This library preparation strategy prefers longer transcripts and was selected against short sRNA transcripts. The strategy was proved to be successful because 2 out of 3 Hfq-bound sRNAs evident by CLIP-Seq lost Hfq-bound signals from the longer transcripts were selected by RIP-Seq library (Fig S3B). We showed that 67.5% intergenic regions and 70.3% antisense regions showed Hfq-bound evidences from CLIP-Seq data, while 49.1% and 30.4% corresponding regions obtained Hfq-bound evidences from a set of RIP-Seq data (Fig S3C). When compared with the CLIP-Seq binding profile, the specifically reduced binding capacity in the intergenic and antisense RNAs but not mRNAs reflected by long transcripts preferred RIP-Seq libraries suggests that the intergenic and antisense RNAs are generally short transcripts. Their higher Hfq-bound efficiency indicates an unexpected global function in gene regulation.

The 5’ leaders are well-known to regulate bacterial gene expression [8]. The regulatory role of the 3’ region of mRNA genes is emerging recently [40]. We explored the Hfq binding profiles in these two classes of non-coding regions in *Y. pestis*. Compared to the 5’ leader regions (Fig 2C, left), we showed that Hfq binding is strongly enriched at the 3’ regions downstream mRNA genes, which was evident both by Hfq CLIP-Seq and RIP-Seq reads (Fig 2C, right). The 3’ binding region was peaked immediately downstream of the stop codons for about 75nt, and extended to over 100nt upstream (Fig 2C). This Hfq-binding profile suggests a much more global regulatory role of the 3’ region in *Y. pestis*.

The mRNA 3’ regions have been reported to encode two major types of sRNAs. Type I is independently transcribed from the 3’ end of a mRNA and type II is processed by an endonuclease at the 3’ region of the mRNA from a primary transcript [40]. The four reported that type I and II sRNAs from *E. coli* and *Salmonella* were examined for their transcripts and Hfq binding profiles in *Y. pestis.* Three of the host mRNA genes were well expressed in *Y. pestis*, including the type I MicL in *cutC* mRNA and type II SroC and CpxQ sRNAs located at the 3’ of *gltl* and *cpxP* mRNA. All of these three 3’ sRNAs were bound by Hfq (Fig 2D and Fig 1D).

### Hfq Binding Sites and Motifs in the Coding and Non-coding Regions

A total of 2511 CLIP-Seq peaks and 1518 RNA-Seq peaks were recovered based on the sequencing data obtained from Hfq-FLAG strain (Table S5). The Hfq-bound regions were peaked around 150nt in length, and could be as long as 500nt and longer (Fig 3A). Such a long Hfq-binding region could be partially resulted from the partial RNase T1 digestion during operation. In contrast, the transcript peaks were quite speeded, and generally longer than Hfq-bound peaks (Fig 3A). In general, Hfq has a larger tendency to associate with the non-coding regions including both the intergenic and antisense (Fig 3B)

**Fig 3.**
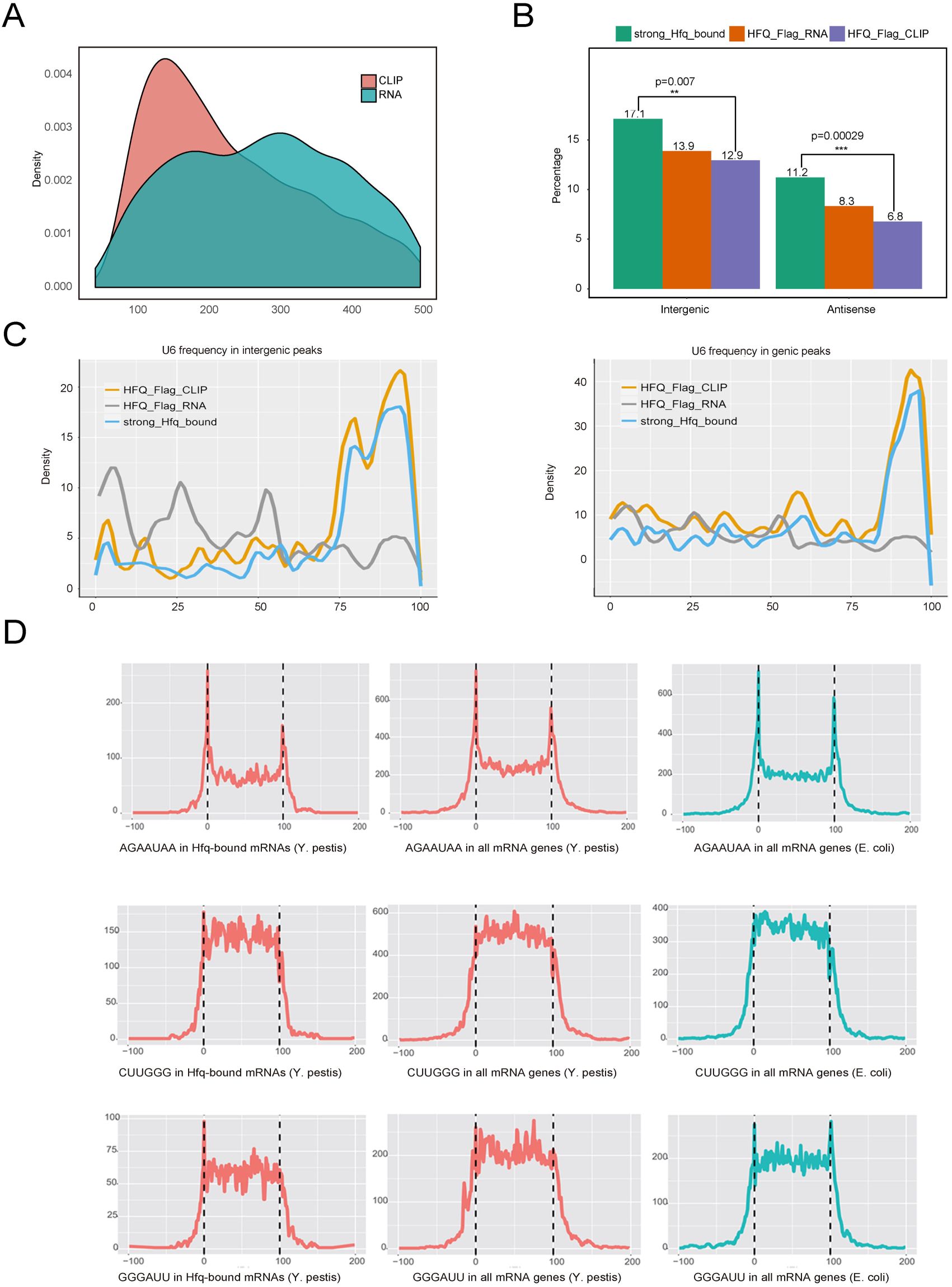
Hfq binding sites and motifs distribution in the coding and non-coding regions. (A) Peak length distribution of CLIP-Seq and RNA-Seq experiments. (B) Intergenic and antisense peak number percentage of RNA-Seq, CLIP-Seq, and strong Hfq-bound peaks. P-value was labeled to illustrate the enrichment of peak distribution in antisense and intergenic region. (C) U_6_-stretch frequency and position in three classes of peaks, including all CLIP-Seq peaks, all RNA-Seq peaks and strong Hfq-bound peaks. Presentation was separated by intergenic (left) and genic (right) regions. (D) Motif distribution along the Hfq-bound mRNAs (left), all mRNAs (middle) of *Y. pestis*, and in all mRNAs of *E.coli* (right). From top to bottom, the figures represent for AGAAUAA, CUUGGG and GGGAUU motifs, respectively.

Theoretically, CLIP-Seq peaks indicated the Hfq-bound regions, while RNA-Seq peaks indicated the steady-level transcripts. The latter is expected to cover the former. We then selected strong Hfq-bound peaks in which CLIP peaks containing 4-fold more CLIP-Seq reads than RNA-Seq reads, resulted in 1499 qualified peaks. Among these, 1168 overlapped the known genes; 131 overlapped the antisense strands, and 200 in the intergenic regions (Fig 3B). The selection criteria are quite strict, as reflected by the loss of four of the five Hfq-bound sRNAs and all three Hfq-bound tRNAs identified above (Fig 1).

Running Homer algorithm [41] recovered highly represented Hfq binding motifs from these three different classes of peaks. These cellular motifs harbor all three known types of motif sequence including polyU, A-rich, and UA-rich bound by the proximal, distal and rim surfaces of Hfq respectively. Hfq-bound RNA motifs in living *Y. pestis* were more conserved in short motif sequence composition but quite flexible in motif organizations. For example, the top motif AAUAA was highly represented in mRNAs, intergenic RNAs and antisense RNAs (Table 1). The two conserved nucleotides preceding this motif were AG(C) in mRNAs, AG in intergenic and UA in antisense sRNAs. The resulted motif composition contained a combination of ARN and UAA motifs in mRNAs and intergenic RNAs, and two UAA motifs in the antisense. Moreover, our results revealed a previously unrecognized G-rich motif. GGGGAUU motif was highly represented in Hfq-bound mRNAs and intergenic sRNAs, but not in antisense sRNAs. G-rich motif might contact Hfq at the distal face as the ARN motif does [26].

**Table 1.**
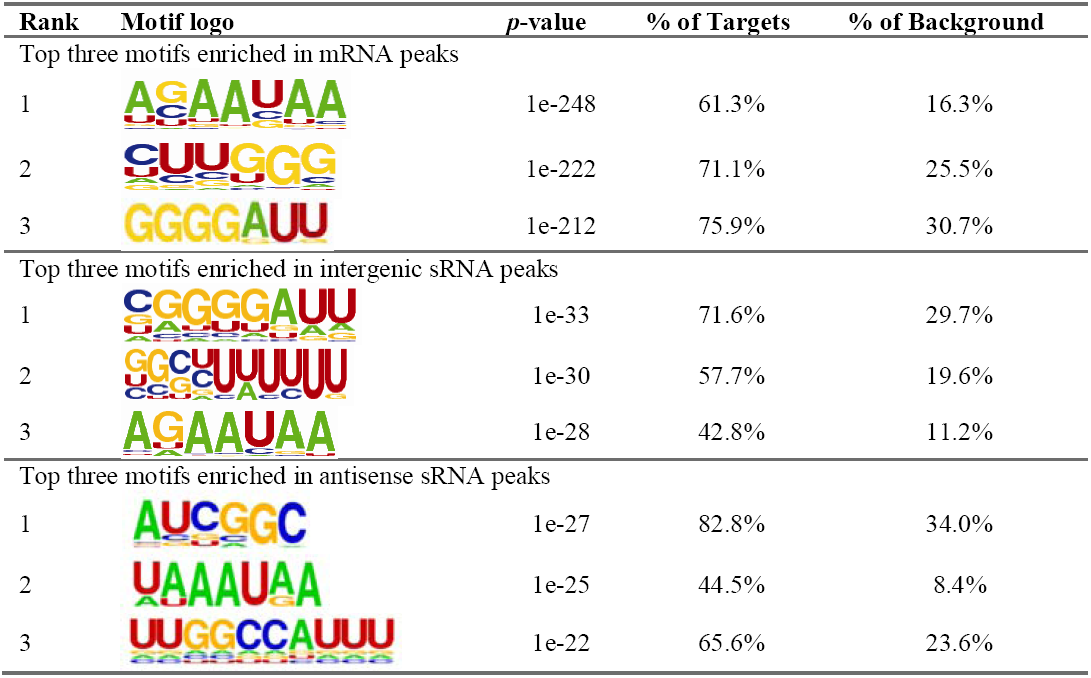
Top three consensus logos generated from mRNAs and sRNAs bound by Hfq

| Rank | Motif logo | *p*-value | % of Targets | % of Background |
| --- | --- | --- | --- | --- |
| Top three motifs enriched in mRNA peaks |  |  |  |  |
| 1 |  | 1e-248 | 61.3% | 16.3% |
| 2 |  | 1e-222 | 71.1% | 25.5% |
| 3 |  | 1e-212 | 75.9% | 30.7% |
| Top three motifs enriched in intergenic sRNA peaks |  |  |  |  |
| 1 |  | 1e-33 | 71.6% | 29.7% |
| 2 |  | 1e-30 | 57.7% | 19.6% |
| 3 |  | 1e-28 | 42.8% | 11.2% |
| Top three motifs enriched in antisense sRNA peaks |  |  |  |  |
| 1 |  | 1e-27 | 82.8% | 34.0% |
| 2 |  | 1e-25 | 44.5% | 8.4% |
| 3 |  | 1e-22 | 65.6% | 23.6% |

As a conserved sequence component of ρ-independent terminator, polyU is a symbol motif of the Hfq-bound sRNAs. We found that the U_6_-stretch motif was presented in 57.7% of the intergenic RNA peaks (Table 1). The number of U_5_-stretch was 2-fold more than U_6_-stretch in the intergenic sRNA sequences bound by Hfq. As expected, the U_6_-stretch preferentially located at the 3’ end of strong Hfq-bound peaks (Fig 3C, left). No such enrichment was observed for RNA-Seq peaks (Fig 3C, left). A population of Hfq-bound mRNA peaks also contain the U_6_-motif at the 3’ end (Fig 3C, right). Such a U_6_-strech enrichment at the 3’ end was not much evident for the antisense RNAs (Fig S4A). It is noteworthy that U_5_-motif occurred at a much higher frequency with a pattern similar to the U_6_-stretch (Fig S4A). The presence of polyU motif at the 3’-termi of Hfq-bound mRNA suggests that Hfq could use its proximal surface to contact with mRNA as well, consistent with the recently identified class of sRNAs located in 3’ regions of mRNAs.

### Motif Complementarity in Intergenic sRNAs and between sRNAs and the Coding Regions

Consistent with their Hfq-bound feature, all classes of Hfq-bound peaks tended to form local secondary structures (Fig S4B). The most striking feature of the top combinatory motifs was the strong complementarity between U_6_-stretch motif in intergenic sRNAs and either AGAAUAA or GGGGAUU presented in sRNAs and mRNAs (Table 1). Statistical analysis showed that U_6_-strech was located close to GGGGAUU and/or to AGAATAA sequence for base-pairing in intergenic sRNAs (Fig S4C). In addition, we analyzed the distribution of the top three combinatory motifs on the mRNAs harboring strong Hfq-bound sites. AG/CAAUAA motif was peaked at the 5’ and 3’ ends of the target mRNAs, while the other two highly represented motifs CUUGGG and GGGAUU were presented in the body regions of mRNAs (Fig 3D, left panels). We wondered whether the motif selection was caused by Hfq selection. Analysis of the sequence composition of mRNAs of both *Y. pestis* and *E. coli* showed that the above motif patterns were true for all mRNAs (Fig 3D, middle and right panels). Therefore, the location specificity of these Hfq-bound mRNA motifs should not be caused by a selection of Hfq binding, it is rather an intrinsic feature of bacterial mRNA structure. Nevertheless, we noticed that all classes of motifs located at the 5’ end were more selected than those located at other regions (Fig 3D, left and middle)

We found that 53.64% and 70.83% mRNAs from *Y. pestis* and *E. coli* respectively contain either an AGAAUAA, CUUGGG or GGGAUU motif. 71.82% of Hfq-bound mRNAs contain at least one of these top motifs. AGAAUAA motifs at the 5’ and 3’ termini were present at a similarly high frequency in both *Y. pestis* and *E. coli*. However, CUUGGG and GGGAUU were present at a much higher frequency in *Y. pestis* than *E. coli*, which could represent species-specificity in Hfq-binding mediated gene regulation.

### Small RNAs Identified by Hfq-bound Reads Differ Greatly from those by RNA-Seq Reads

We wanted to identify *Y. pestis* sRNAs to further understand the binding features of Hfq-sRNA by using the RNA-Seq data obtained from the same bacterial strains and similar culture condition as for generating CLIP-Seq data. Although previous studies have identified hundreds of *Yersinia* sRNAs, these sRNAs are not well correlated among different studies, even for the same species *Y. pestis* [42–45] (Fig S5A).

We predicted that sRNAs from RNA-Seq reads mapped in the intergenic regions and antisense regions of *Y. pestis* 91001 genome using the transcript peak algorithm described above. The amount of transcription peaks (antisense and intergenic) in Hfq-FLAG strain (ΔHfq background), FLAG tag strain (Hfq wild-type background), and Hfq wild-type strain without any plasmid were 246, 313 and 237 respectively. These peaks were strongly overlapped with each other, and 178 of them share over 80% overlapped sequence, which were considered as the same sRNAs (Fig 4A). After merging, 373 sRNA transcripts were identified from all three strains. Over 72% were ranged from 70nt to 260nt in length (Fig 4B). Among the intergenic sRNAs, only 40 of them harbor a canonical terminator within 150nt of their 3’ ends, while 85 harbored a canonical promoter within 150nt of their 5’ end (Fig S5B). Among them, 12 harbor both the terminator and promoter. Over 40% of the previously identified *Yersinia* sRNAs from different studies found their homologs among these 373 sRNAs (Table S5).

**Fig 4.**
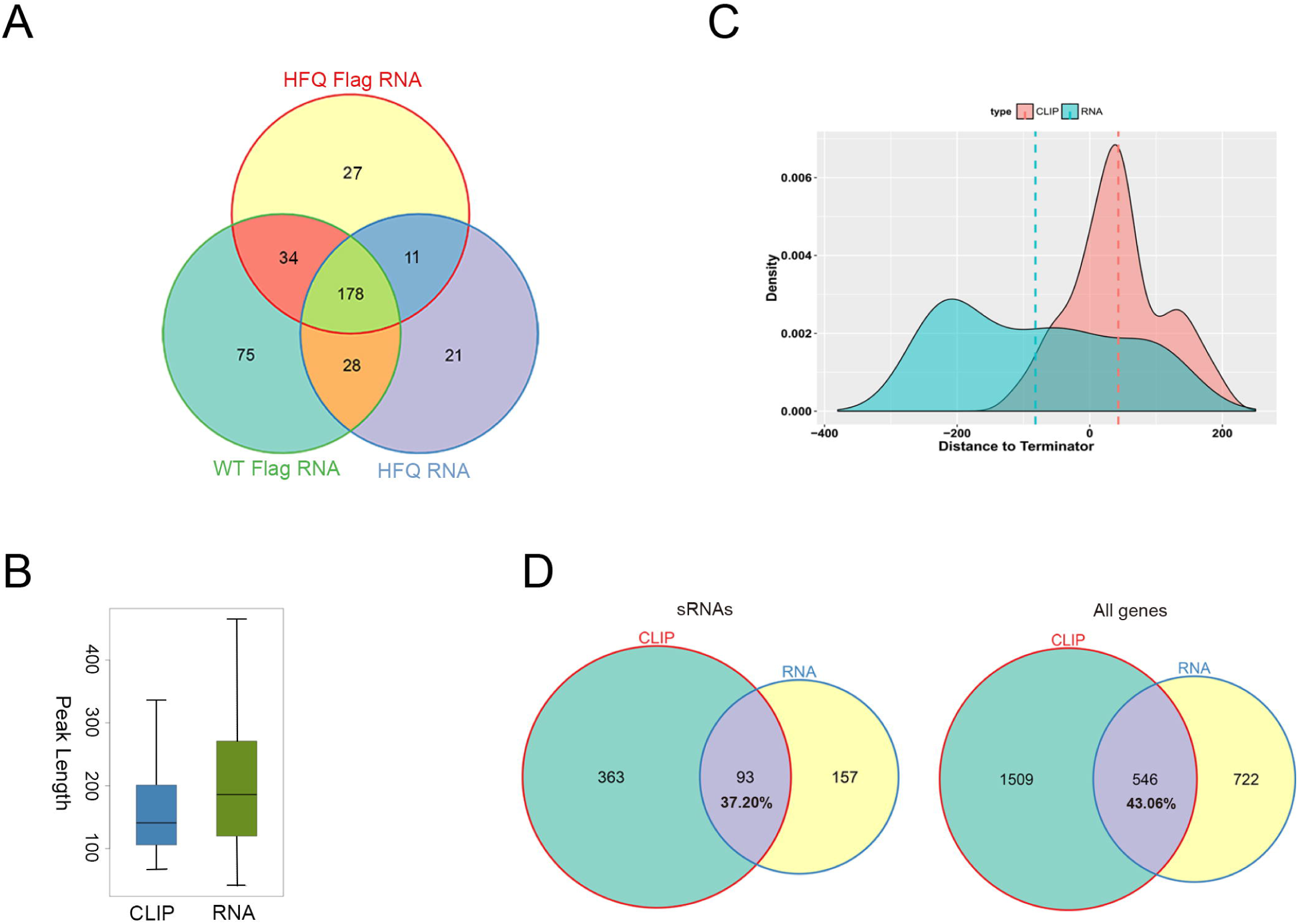
Candidate sRNAs identified by RNA-Seq and CLIP-Seq. (A) Venn diagram showing the overlap of sRNA predicted from RNA-Seq data of three different *Y. pestis* strains. (B) Box plot of the length of sRNA peaks obtained from CLIP-Seq and RNA-Seq data. (C) Density plot of the distance distribution of peak end to terminator. Dashed line represents for the mean distance to terminator. (D) Venn diagram analysis of the overlap between Hfq-bound peaks (CLIP-Seq, left) and overall transcript peaks (RNA-Seq, right).

Analysis of Hfq-bound sRNA peaks from the CLIP-Seq reads revealed 456 qualified intergenic and antisense peaks (Table S6). Hfq-bound sRNAs were shorter than the RNA-Seq sRNAs (Fig 4B). Compared to the RNA-Seq sRNA peaks, Hfq-bound sRNA peaks were closer to canonical transcription terminators and more distant to the promoters (Fig S5B). Most of Hfq-bound sRNA peaks were located downstream of the predicted terminators (Fig 4C). When we set one nucleotide overlap as a criterion, about 2/3 of Hfq-bound sRNA peaks were not overlapped with the sRNA transcript peaks, similar to Hfq-bound mRNA peaks (Fig 4D). Such a low overlap efficiency was unexpected considered the high coverage of cDNA libraries. The low overlap efficiency of both sRNA and mRNA peaks between CLIP-Seq and RNA-Seq data ruled out the possible inefficiency of RNA-Seq libraries in recovering small transcripts. The more reasonable explanation could be Hfq-binding induced destabilization of sRNAs and mRNAs.

### RNA Segments Downstream of Hfq-bound Sites in both sRNAs and mRNAs Were Destabilizaed

To further explore the above hypothesis, we separated sRNA peaks into three different classes. Stable Hfq-unbound sRNA peaks (type I) refer to those having RNA-Seq transcript peaks only, including 156 sRNA members. Unstable Hfq-bound sRNA peaks (type II) refer to those having Hfq-bound peaks only, including 361 members. Stable Hfq-bound sRNA peaks (type III) refer to those with both Hfq-bound and transcript peaks being overlapped at least one nucleotide, including 93 members (Table S7). The stability of these three classes of sRNA peaks was reflected by their transcript abundance recovered from RNA-Seq data (Fig 5A). We plotted the length distribution of these three classes of peaks, showing the overlapped peaks were generally longer than the non-overlapped peaks (Fig S6A).

**Fig 5.**
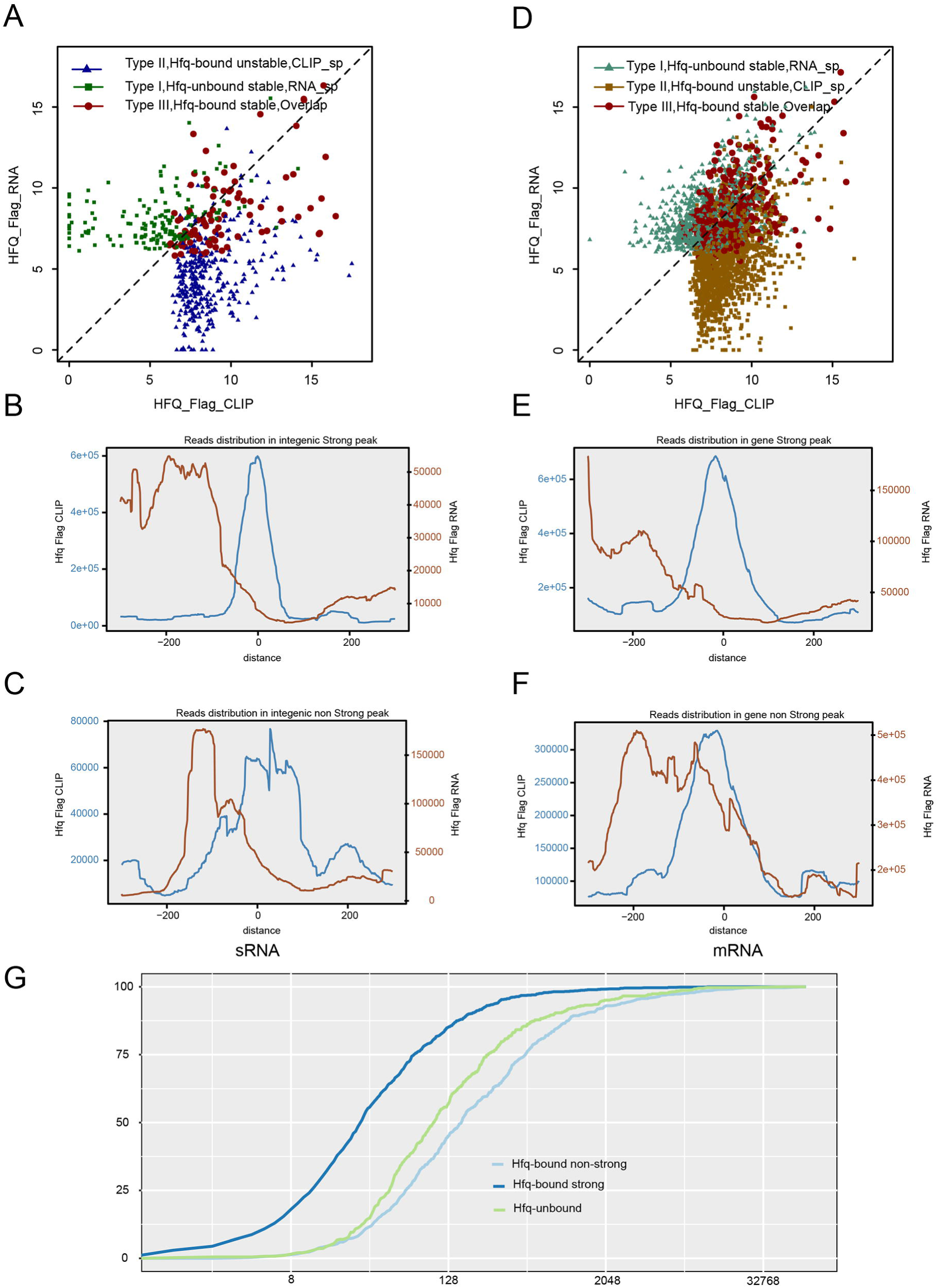
Hfq-bound sRNA and mRNA segments were generally unstable. (A) Dot plot of the abundance of three classes of sRNA peaks: Hfq-unbound stable, Hfq-bound stable, and Hfq-bound unstable. (B) Distribution profiles of all CLIP-Seq and RNA-Seq reads around the center of Hfq-bound strong intergenic peaks. (C) Distribution profiles of all CLIP-Seq and RNA-Seq reads around the center of Hfq-bound non-strong intergenic peaks. (D) Dot plot of the abundance of three classes of mRNA peaks: Hfq-unbound stable, Hfq-bound stable, and Hfq-bound unstable. (E) Distribution profiles of all CLIP-Seq and RNA-Seq reads around the center of Hfq-bound strong mRNA peaks. (F) Distribution profiles of all CLIP-Seq and RNA-Seq reads around the center of Hfq-bound non-strong mRNA peaks. (G) Accumulative plot of gene expression abundance. Genes were divided into three groups by Hfq binding: Hfq-bound non-strong peak genes, Hfq-bound strong peak genes, and Hfq-unbound genes.

We then plotted CLIP-Seq and RNA-Seq reads around the center of strong Hfq-bound peaks in sRNAs to study the RNA abundance around Hfq-bound sites. Interestingly, the distribution of RNA-Seq reads inside of and downstream of Hfq-bound sites strongly declined compared with that of the upstream (Fig 5B), which supported the hypothesis of Hfq-induced destabilization of sRNA segments downstream of the Hfq-bound sites. The strong peaks selected against of peaks with high-abundance of nearby transcript reads. We then plotted CLIP-Seq and RNA-Seq reads around the center of Hfq-bound sites recovered from non-strong intergenic CLIP peaks, a similar declined abundance of RNA-Seq reads downstream of the Hfq-bound sites was observed (Fig 5C). Interestingly, a highly-abundant transcript peak upstream of the Hfq-binding center was observed for non-strong intergenic CLIP peaks, showing a distance of about 120nt (Fig 5C).

The distribution of Hfq-bound cDNA reads and transcript cDNA reads in individual sRNA peaks were plotted, showing examples of three classes (Fig S6B). We also showed the stability of these sRNAs in response to Hfq deletion. Northern blot analysis of sRNAs in the wild-type *Y. pestis* strain and the ΔHfq strain (Fig S6B) showed that the knock-out of Hfq decreased the stability of almost all Hfq-bound sRNAs, regardless of their differential stability in the Hfq+ strain. In contrast, the abundance of all Hfq-unbound sRNAs was not affected by Hfq deletion (Fig S6B). These results are consistent with a model that the destabilization of Hfq-bound sRNA segments depends on their base-pairing with target mRNAs facilitated by Hfq binding [16]. As a result, the loss of Hfq also removes the mRNA target-dependent destabilization of Hfq-bound sRNAs.

The strong correlation between Hfq-binding and sRNA destabilization stimulated us to propose that the stable Hfq-unbound sRNA may lack Hfq-binding motifs. Analysis of the overrepresented motifs in all 373 sRNA peaks identified from RNA-Seq revealed the lack of typical Hfq binding motif in sRNAs, U_6_-stretch (Fig S6C).

Destabilization of Hfq-bound sRNAs could be resulted from coupled degradation of a small regulatory RNA and its mRNA targets [16]. The differential transcript abundance of three classes of mRNA peaks were similar to those of sRNAs (Fig 5D). We then analyzed the distribution of transcript reads around the center of Hfq-bound sites from strong and non-strong CLIP peaks recovered from mRNA regions. Strong Hfq binding correlated with the destabilization of the downstream mRNA segments, highly similar to that of sRNAs (Fig 5E). RNA-Seq reads distribution upstream of non-strong Hfq binding sites was almost the same as that of sRNAs, and the downstream destabilization was also evident (Fig 5F). However, the appearance of an RNA-Seq read peak highly overlapped with the CLIP-Seq read peak was quite unique (Fig 5F).

In light of the proposed mechanism of Hfq facilitated sRNA-mRNA degradation, we explored the relationship between Hfq binding of mRNA and their stability. The accumulative abundance of mRNAs harboring strong or non-strong Hfq peaks, was plotted, showing that mRNAs containing non-strong peak genes were more abundant than those containing strong peaks (Fig 5G). Genes containing Hfq-unbound transcripts peaks were generally expressed at levels significantly lower than those containing the Hfq-bound (Fig 5G).

### Hfq Regulates the Processing of Hfq-bound sRNAs

Several Hfq-binding sRNAs including GlmZ and SraH/ArcZ are reported to experience the RNase E processing, while GcvB contains two transcriptional termination sites to produce two sRNA isoforms [46–50]. The proposed secondary structure of *Y. pestis* GlmZ is almost identical to that of *E. coli* (Fig 6A), while that of SraH/ArcZ was strikingly different (Fig 6B) [50]. The two sRNA isoforms for both GlmZ and SraH/ArcZ were detected by Nornther blot analysis, while the knock-out of Hfq has opposite effect on their processing (Fig 6C-D). The RNase E processing sites on GlmZ and SraH/ArcZ were readily recognized from the Hfq binding profiles. The processing site of SraH/ArcZ was exactly the same as previously reported from *E. coli* [50]. However, the processing site of GlmZ was different from the previous report site (Fig 6A). In both cases, the processing resulted in a U-rich 5’-end, which can access the A-rich sequence enriched at the 5’ and 3’ ends of mRNAs (Figs 6A-B). Interestingly, *Y. pesti* GcvB does not display two transcript isoforms, although its sequence is highly similar to *E. coli* (Fig 6E). Ffs was a negative control for Hfq binding. Hfq binding-regulation of sRNA processing seems to be global (Fig 6F), because among other three sR128, sR142 and sR132 showing distinct isoforms, the isoform ratios of two were changed upon Hfq deletion (Fig S6B).

**Fig 6.**
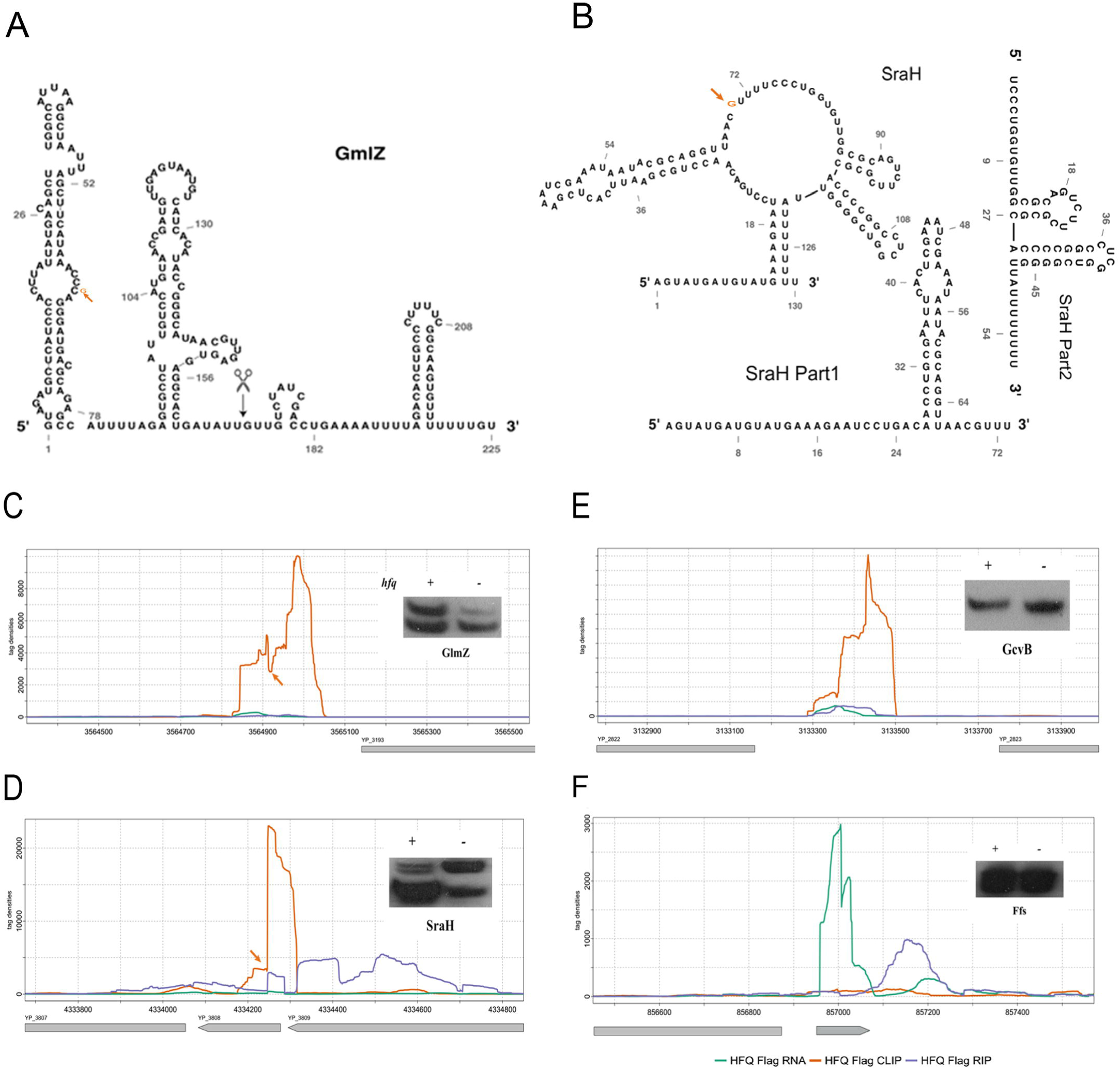
Hfq Generally Regulates the Processing of Hfq-bound sRNAs. (A) Second structure of the *Y. pestis GlmZ* sRNA. The position of red arrow represents for the processing sites by Hfq binding. The black scissor represents for the reported processing site from *E. coli*. (B) Second structure of the *Y. pestis SraH* sRNA and the two parts after processing. The position of orange arrow represents for the processing sites by Hfq binding. SraH Part1 and Part2 represents for the second structure after processing. (C-F) Reads distribution and Northern blot of four sRNAs. The red arrow points the RNase E cleavage sites, corresponding to the shift site of Hfq-binding peaks. *GcvB* and *Ffs* show no processing sites. (A) Please be noted that *Ffs* sRNA presented here was identified as a sRNA peak identified from RNA-seq data (Table S7), which was on the opposite direction of the one originally annotated in the genome (Table S2, S3 Fig).

## DISCUSSION

Sm proteins are a family of small proteins that assemble the core components of the U1, U2, U4 and U5 spliceosomal snRNPs, and therefore are central for eukaryotic pre-mRNA splicing [51]. Lsm proteins containing the ‘Sm motif’ are then discovered to function in eukaryotic mRNA decaping and decay. Lsm1 to Lsm7 proteins interact with both the mRNA and the mRNA degradative machinery to exert the function [52, 53]. Sm-Lsm protein family is presented in all three domains of life. Hfq has been known for over a half century and is a typical LSm protein [3]. By cooperation with diverse sRNAs, Hfq has been shown to play a key role in degrading bacterial mRNAs. The process involves in the recruitment of RNase E, a key member of RNA degradosome [7, 14]. Decay of mRNA can be either coupled with sRNA or not [4, 5, 54]. Hfq is known to use its proximal, distal and rim surfaces to contact both sRNA and mRNA with specific nucleotide preference, and therefore promoting sRNA-mRNA base-pairing [10]. There are several fundamental questions waiting to be addressed in living bacteria, including: (1) how many sRNAs and mRNAs are contacted by Hfq? (2) how do the different Hfq surfaces contact sRNA and mRNA in living cells? (3) how does the Hfq binding contribute to sRNA and mRNA base-pairing and their decay in bacterial cells? In this study, we obtained Hfq-bound RNA using both CLIP-Seq and RIP-Seq techniques. By setting RNA-Seq data as controls and developing proper algorithms to analyze the high-quality data, we were able to address these three questions to a good depth and propose a model for Hfq binding and facilitation of sRNA-mRNA coupled degradation in living bacteria.

### Hfq Extensively Binds mRNAs

Although several co-immunoprecipitation studies have revealed that Hfq binds tens of sRNAs in both *E. coli* and *Salmonella* [36, 37, 55, 56] and hundreds of mRNAs in the latter organism [37], more information on Hfq-bound RNA targets in other bacteria is valuable to better understand Hfq action in living cells. By expressing Hfq-FLAG protein in a Hfq knock-out background, we used anti-FLAG antibody to obtain one set of high-quality CLIP-Seq data and two sets of high-quality RIP-Seq data. Using Hfq-bound 23S rRNA as control, we ambiguously identified that thousands of expressed mRNA genes (∼80%) were bound by Hfq. These results lead to a hypothesis that Hfq might control the stability of most mRNAs with its sRNA partners.

A more recent study of Hfq-bound sRNAs of *Salmonella typhimurium* under different growth stages reveals that 3’ UTRs represent a genomic reservoir of regulatory small RNAs [57]. Here we revealed a much larger number of such sRNAs in *Y. pestis*.

### Hfq Flexibly Contacts sRNA and mRNAs Using Multiple Surfaces: Formation of Hfq-sRNA-mRNA Complex

*In vitro* studies have revealed that Hfq uses its proximal face to bind polyU sequence in sRNAs, and its distal and rim surfaces to contact with A-rich and UA-rich sequences in sRNA and mRNAs [10]. Hfq-bound RNA motifs from our CLIP-Seq data revealed a more comprehensive Hfq binding strategy in living cells. In the mRNAs, in addition to A-rich and UA-rich motifs, Hfq-bound G-rich and UG-rich motifs have been identified in *Y. pestis* cells. These motifs mirror A-rich and UA-rich sequences and may be contacted by the distal and rim surfaces of Hfq. In the cases of sRNAs, the top three motifs were featured either by the canonical terminator sequence containing U_6_-stretch motif proceeded by GC-rich sequence (Table 1), or by other two motifs that are G-rich or A-rich. U_6_-stretch motif can extensively base-pair with either G-rich or A-rich motifs.

The *in vivo* Hfq motifs are comprised of different known short motifs, enabling an Hfq hexamer to use different surfaces to recognize and effectively contact a RNA sequence. The combinatory organization of different motif blocks could allow a specific RNA sequence to contact multiple faces of an Hfq hexamer. This organization could have additional advantages in assembly of Hfq-RNA complex.

The findings presented expanded our understanding of the dynamic and efficiency of Hfq in binding mRNAs. Hfq is capable of associating with newly transcribed mRNAs and sRNAs with a low affinity. However, its binding of mRNA is effectively excluded by the transcription-coupled translation and its binding of sRNAs is stabilized by the presence of nearby secondary structure. The Hfq-sRNA complex interacts with the translationally inactive and/or repressed mRNAs, which enables the formation of intermolecular base-pairing between sRNA-mRNAs and the increase of local concentrations of RNase E for cleavage of sRNA-mRNA duplex (S6 Fig).

Increasing numbers of sRNAs have been proved to simultaneously act on multiple mRNAs. And likewise, many mRNA transcripts are emerging as shared targets of multiple cognate sRNAs. Since Hfq is assumed insufficient relative to RNA species, RNA is shown to actively cycle by competition for the access to Hfq [9, 58]. The results present in this study suggest that the Hfq-sRNA complexes could select their target in a relative very flexible way. For example, the ubiquitous U_6_ stretch of many sRNAs can base-pair with many mRNAs containing A-rich or G-rich motifs at the terminal parts or the body regions. If this is true, the dynamic and amplitude of Hfq-sRNA-mRNA regulation will be beyond our imagination.

### Hfq-bound sRNAs were Generally Unstable: A Comprehensive List of *Y. pestis* sRNAs

We have demonstrated that Hfq binding of sRNAs is more complex than expected. Firstly, the sRNAs predicted from sequencing the transcript reads from bacterial cells can only represent a population of sRNAs but not all. Secondly, a large population of sRNAs is unbalanced in their stability, with 5’ portion being more stable than the 3’ portion, largely due to the Hfq binding. Therefore, some sRNAs predicted from transcriptome reads may be shorter than the full transcripts. At last, Hfq-binding is associated with sRNA degradation during the normal growth condition where transcription is active.

In summary, sRNAs are highly dynamic in their transcription and degradation. Identification of full length sRNA genes is challenging. The challenge is further complicated by the lack of canonical terminators in many sRNAs, and the presence of sRNAs overlapping the 3’ end of mRNA genes [38, 40].

In this study, we generated a comprehensive sRNA list comprising about 700 members encoded by a vaccine *Y. pestis* genome, only a small fraction of them have been identified before. This list does not include those overlapped the 3’ region of mRNA genes. About 363 Hfq-bound sRNAs difficult to be identified by transcriptome sequencing were identified. We therefore established a concept that Hfq binds hundreds of sRNAs, which could be involved in controlling the stability of most if not all of mRNAs in a bacterial species.

### Hfq-bound Sites Define Two Positions for the Couple-degradation of the Base-paired sRNA-mRNA: A General Mechanism for Cellular mRNA Surveillance

The mechanism of coupled degradation of sRNA-mRNA complex is feasible for a quick response of environmental change by bacterial cells [7], and has been proposed in 2003 [16] but has few further direct evidences afterward [4, 7, 14]. In this study, genome-wide analysis of multiple classes of sRNAs and mRNAs in their aspects of the Hfq-bound capability allows us to comprehensively revisit this issue. We have demonstrated that sRNA segments at the Hfq-bound sites and their downstream are globally unstable. The mRNAs containing strong Hfq-bound peaks showed the similar pattern as Hfq-bound sRNA did. However, mRNAs containing non-strong Hfq-bound peaks showed that the Hfq-binding sites are protected, while the downstream segments are destabilized. In light of the RNase E function in Hfq-sRNA mediated RNA degradation [14], we proposed that Hfq-bound sites render two positions for the RNase E entry, which will result in the degradation of mRNA segments downstream of Hfq binding sites but different decay fates of the Hfq-bound sites, via a 5’P-dependent RNase E degradation pathway (S7 Fig). The coupled degradation of both mRNA and sRNA in the RNase E-bound sites lead to either the direct degradation of the 5’-P containing sRNAs or recycling of broken sRNAs containing either 5’-P or 5’-PPP. Recycling of broken sRNAs explains the lack of protected Hfq-bound sites in sRNAs.

Interestingly, we showed that Hfq-bound AGAAUAA motifs are located at both the 5’ and 3’ termini of *Y. pestis* and *E. coli* mRNAs sites, while CUUGGG and GGGAUU are located at the body regions of the genes. All these sequence motifs are partially complementary to U-rich sequence in sRNAs. Given the large diversity of sRNA sequences, it is not surprising that Hfq-sRNA has a chance to bind most of cellular mRNAs and to mediate their degradation when they are not effectively translated. Although there are Hfq-bound mRNA motifs, we find that Hfq binding of mRNAs lacks selectivity because the motifs were features of mRNA gene sequence. The lack of selection of the binding site endorses the RNA chaperon Hfq to surveil the RNA hemostasis of the whole bacterial transcripts via the cooperation with its partner sRNAs.

## MATERIAL AND METHODS

### Construction of C-terminally Flag-tagged plasmids

By using a fusion PCR protocol, oligonucleiotides encoding 3 × Flag affinity tag (DYKDHDGDYKDHDIDYKDDDDK) were added before TAA termination codon of the Hfq gene. A fragment covering a region of 298 nt upstream, the entire *hfq* gene followed by 3 × Flag and 176 nt downstream was cloned into the multiple cloning site of plasmid pACYC184, designated as pHfq-Flag. The other fragment spanning a region of 298 nt upstream, the first 21 nt of the *hfq* gene followed by 3 × Flag and 176 nt downstream was also introduced into pACYC184, designated as Flag.

### Bacterial strains and growth conditions

*Y. pestis* wild-type strain 201 belongs to a newly established *Y. pestis* biovar, microtus, which is avirulent in humans but highly lethal in mice. The *hfq* deletion strain Δ*hfq* was generated by λ-Red homologous recombination methods as previously described [35]. The *Y. pestis* Hfq-Flag and WT-Flag strains were constructed by transforming the pHfq-Flag and Flag mentioned above into Δ*hfq* and the WT strain, respectively. Bacteria were grown in brain heart infusion (BHI) broth (Difco) supplemented with appropriate antibiotics overnight at 26 °C with shaking at 200 rpm until exponential growth phase (OD_620_ = 0.8). Bacterial growth was stopped by centrifugation for 6 min at 5,000 rpm at 4 °C. The pellets were frozen into liquid nitrogen and stored at −80 °C until cells were lysed. Western blotting was performed by using monoclonal FLAG antibody (Sigma) to detect the Flag-tagged proteins those were present within cells of Hfq-Flag and WT-Flag.

### RNA-seq, CLIP-seq and RIP-Seq

For RNA-seq, total RNAs were extracted from *Y. pestis* Hfq-Flag and WT-Flag strains mentioned above by using Trizol Reagent (Invitrogen). For CLIP-seq, the cultures grown under the same conditions were collected and resuspended in 10 mM Tris-HCl (pH8.0), respectively. The pellets were dispersed on a petri dish and irritated uncovered with 400 mJ/cm^2^ of UV 254 nm to form the crosslinked RNA-protein complex. Bacterial cells were collected and lysed in RIP Lysis buffer and subjected to co-immunoprecipitation (Co-IP). Co-IP was carried out to isolate the FLAG-bound and the Hfq-Flag-bound RNA by using Flag antibody according to the manufacturer’s instructions of RNA-Binding Protein Immunoprecipitation Kit (MILLIPORE). Briefly, the lysate was centrifuged at 12,000g at 4°C for 10 min. The clear lysate was incubated with 1.0 mL of beads-antibody complex in RIP Immunoprecipitation Buffer, followed by incubation at 4 °C for 3 h on a rotator. Briefly centrifuge the immunoprecipitation tubes and place on the magnetic separator and discard the supernatant. The anti-FLAG beads were then washed total six times with 0.5 mL of RIP Wash Buffer and digested by RNase T. The immunoprecipitated RNA fragments were radiolabeled using PNK and separated by SDS-PAGE gel. The bands corresponding to the equivalent size of Hfq protein were cut out and purified. The crosslinked RNA-protein complexes were digested with proteinase K at 55 °C for 30 min. RNA was extracted using Trizol and phenol:chloroform, followed by isopropanol precipitation. The purified RNA was treated with DNase I (Promega) and sequenced using the Illumina/Solexa RNA-sequencing protocol.

For RIP-Seq, 500 μL lysate was incubated with 10 ug anti-Flag antibody or control IgG antibody overnight at 4 oC. The immunuprecipitates were further incubated with protein A Dynabeads for 1h at 4 oC. After applying to magnet and removing the supernatants, the beads were sequentially washed with lysis buffer, high-salt buffer (250 mM Tris 7.4, 750 mM NaCl, 10 mM EDTA, 0.1% SDS, 0.5% NP-40 and 0.5 deoxycholate), and PNK buffer (50 mM Tris, 20 mM EGTA and 0.5% NP-40) for two times, respectively. The immunoprecipitates were eluted from the beads with elution buffer (50 nM Tris 8.0, 10 mM EDTA and 1% SDS) and the RNA was purified with Trizol reagent (Life technologies).

Purified RNAs were iron fragmented at 95oC followed by end repair and 5’ adaptor ligation. Then reverse transcription was performed with RT primer harboring 3’ adaptor sequence and randomized hexamer. The cDNAs were purified and amplified and PCR products corresponding to 200-500 bps were purified, quantified and stored at −80 °C until used for sequencing.

For high-throughput sequencing, the libraries were prepared following the manufacturer’s instructions and applied to Illumina GAIIx system for 80 single-end sequencing by ABlife. Inc (Wuhan, China).

### CLIP-Seq and RIP-Seq raw data clean and alignment statistics

Raw reads were first discarded if containing more than 2-N bases. The reads were then processed by clipping adaptor and removing low quality bases (less than 20). Too short reads (less than 13nt) were also dropped. FASTX-Toolkit (Version 0.0.13) was used to get the clean reads. After that, clean reads were aligned to the *Y.pestis* 91001 reference genome [59] by bowtie2 [60] with no more than 1 seed mismatch. Aligned reads with more than one genome locations were discarded due to their ambiguous locus. Uniquely localized reads were used to do the following analysis. Other statistical results such as gene coverage and depth, reads distribution around start codon and stop codon, were also obtained.

### RNA-Seq raw reads data clean and alignment statitstics

RNA-Seq raw reads were processed by the same pipeline of CLIP-Seq and RIP-Seq except that reads less than 16nt were dropped. After alignment, RPKM value of each gene was obtained from the uniquely aligned reads. To have an overview of the influenced genes by Hfq, we took the differentially expressed genes (DEG) analysis between samples. The DEG groups were Hfq flag vs WT flag, WT flag vs WT. We treated genes with fold change no less than 2 and p-value no more than 0.01 as DEGs. The DEG analysis was performed by edgeR [61].

### Peak Calling method and sRNA definition

To have an exact prediction of *Y. pestis* sRNA and Hfq binding site, we realized an algorithm to detect peaks from alignment results among intragenic, intergenic and antisense regions. 5 bp window size was chosen as the default window size. Peak starting site was identified as the end of one window, the median depth of whichis no more than 0.25 fold of all of the adjacent following eight windows. Peak terminal site was identified as the start of one window whose median depth is no more than 0.25 fold of all of the adjacent previous eight windows. After the algorithm realization, we then filtered the peaks according tothe following three thresholds: 1) the length of peaks should range from 40bp to 500bp; 2) the maximum height of one peak should be no less than 60; 3) the medium height of one peak should be no less than 20 nt. After peak definition, we classified the peaks into three different classes according to their locations: 1) intragenic peaks were defined as peaks whose locus were overlapped with known mRNA genes on the same strand; 2) antisense peaks were defined as peaks whose locus were overlapped with known mRNA genes on the opposite strand; 3) intergenic peaks were defined as neither intragenic nor antisense peaks. Antisense and intergenic peaks were defined as sRNAs both in RNA-Seq and CLIP-Seq samples. We performed the peak calling method with CLIP-Seq and RNA-Seq data sets. According to the peak location, we classified peaks into three types: Stable Hfq-unbound peaks (type I) refer to those having RNA-Seq transcript peaks only. Unstable Hfq-bound peaks (type II) refer to those having Hfq-bound peaks only. Stable Hfq-bound peaks (type III) refer to those with both Hfq-bound and transcript peaks being overlapped at least one nucleotide, including 93 members (S7 Table).

### Hfq bound strong peak definition

The strategy of identifying strongly Hfq-bound and -unbound mRNAs and sRNAs was described as follows. Briefly, peaks identified from the above strategy were the candidate bound and unbound site from Hfq Flag CLIP and Hfq Flag RNA samples. The total number of mapped reads within each peak was calculated. The sequencing data were displayed with the relative depth of mapped reads at each position of nucleotides on the global-genome scale. A 4-fold threshold of total base in each peak between CLIP-seq and RNA-seq was used to define the Hfq-bound or –unbound peaks. The corresponding peak genes were defined as the Hfq-bound or Hfq-unbound genes.

### Motif search and distribution

To illustrate the binding nucleotide pattern of Hfq, we searched the RNA motif enrichment by Homer software [41]. Then we realigned the top motifs to the peak sequence by fuzznuc, and motif distribution was plotted by the normalization length of peak.

### Northern Blot

A DIG Northern Starter Kit (Roche) was used to perform Northern Blot according to the manufacturer’s protocol as previously described [62]. Total RNA samples (3 μg) were denatured at 70 °C for 5 min, separated on 6% polyacrylamide-7 M urea gel, and transferred onto Hybond N+ membranes (GE) by electroblotting. The membranes were UV cross-linked and pre-hybridized for 1 hr. RNA probes labeled with DIG-11-UTP by *in vitro* transcription using T7 RNA polymerase were added. The membranes were then hybridized overnight at 68 °C in a DIG Easy Hyb according to the manufacturer’s protocols. RNA was immunologically detected and exposed to X-ray film.

### RIP-PCR

Based on the immunoprecipitation, purified RNAs were thermal treatment at 65 °C for 5min. Then reverse transcription was performed with RT primer harboring 3’ adaptor sequence and randomized hexamer. The cDNAs were amplified by 2X Dream Taq Mix(Thermo), PCR products electrophoresis analysis by agarose gel.

### Functional enrichment analysis

Lists of binding genes were submitted to DAVID [63, 64] web server (http://david.abcc.ncifcrf.gov/) for enrichment analysis. Enrichment clusters were sorted by the enrichment score in descending order. Categories of one cluster were sorted by p-value in descending order. Fold Enrichment, Bonferroni and Benjamini corrected p-value and FDR were also presented for each category of each cluster.

### Other statistical analysis

Hypergeometric distribution test was used to define the enrichment of each GO term and KEGG pathway. Mann-Whitney test was used to define the differential degree of two independent data sets. Other statistical results were obtained by R software.

## Supporting information

## ACCESSION NUMBERS

The accession number for RNA-Seq, CLIP-Seq and RIP-Seq data reported in this paper is NCBI GEO: GSE77555. Reviewer access link is https://www.ncbi.nlm.nih.gov/geo/query/acc.cgi?token=slchiskottabvcr&acc=GSE77555.

## Supporting information captions

**Fig S1. Experimental procedure and Hfq-bound gene profile of CLIP-Seq.** (A) PAGE gel electrophoresis of cellular Hfq under non-denature condition followed by Western blot analysis. (B) Gene expression correlation analysis between Hfq_Flag and WT_Flag by Pearson correlation. “R” represents for correlation coefficient. (C) Western blot of Hfq-RNA complex separated by SDS-PAGE by using anti-Flag monoclonal antibody (left), and electrophoresis visualization of PCR products amplified from cDNA before (middle) and after (right) gel-cutting purification. (D) Functional clusters of Hfq-bound and unbound genes under in vitro vegetative growth condition.

**Fig S2. Experimental procedure of CLIP-Seq and RNA-Seq.**

**Fig S3. Sample correlation and sRNA prediction of CLIP-Seq and RNA-Seq data.** (A) Three-dimensional correlation of the reads density mapped onto all Y. pestis genes from CLIP-seq, RIP-seq and RNA-seq experiments. (B) Reads density illustration of *Y. pestis* known sRNAs in CLIP-Seq, RIP-Seq and RNA-Seq experiments, including *Spf*, *SsrS*, *CsrB*, *Ffs*, *MicF*, and *RnpB*. Plus or minus sign behind experiment represents for the detection state of sRNA in each experiment. Please be noted that the *Ffs* sRNA was from the genome annotation, not the predicted *Ffs* sRNA. (C) Hfq binding of the intergenic and antisense regions. Bar plot presentation of the results from analysis of CLIP-seq and RIP-seq 2nd data.

**Please be noted the *Ffs* sRNA read distribution shown here was according to the previously published genome annotation, which was shown to be wrong in the strand direction. The correct *Ffs* sRNA on the opposite direction was among the sRNA peaks identified from RNA-seq data (S7 Table) and shown to be Hfq-unbound (Fig 6F).

**Fig S4. Motif distribution profile and the analysis of sRNA second structure potential.** (A) U6-stretch frequency and position in antisense peaks of three samples, including CLIP-Seq, RIP-Seq, and RNA-Seq (top). U5-stretch frequency and position in peaks of three samples, including CLIP-Seq, RIP-Seq, and RNA-Seq. Presentation was separated by intergenic, antisense and genic regions from top to bottom. (B) Violin plot of the base-pairing potential of intergenic sRNAs, mRNA and antisense sRNAs. Y label represents the minima free energy. (C) Venn diagram of the overlap among intergenic peaks that three different motifs were found.

**Fig S5. sRNA annotation by homology and structure analysis.** (A) Venn diagram of the overlap among three different stydies (Beauregard et al., 2013; Koo et al., 2011; Saadeh et al., 2015; Yan et al., 2013). (B) Promoter and terminator prediction of sRNAs from CLIP-Seq and RNA-Seq data. sRNAs were classified into intergenic and antisense sRNAs.

**Fig S6. Characterization of sRNAs bound by Hfq.** (A) Violin plot of the length distribution of CLIP-Seq and RNA-Seq peaks. Peaks were divided into three classes: CLIP-specific, RNA-specific, and Overlap. (B) Reads distribution and Northern blot of three type sRNAs classified by Hfq binding. Top left three panels represent Hfq-unbound stable sRNAs (Typle I), Top right three panels represent Hfq-bound unstable sRNAs (Typle II), and the bottom two represent Hfq-bound stable sRNAs (Type III). Please be noted that Northern blot analysis of sRNA level was performed on total RNAs from the Hfq+ (WT) and Hfq- (Hfq deleted) strains. (C) Enrichment of intergenic (left) and antisense (right) Hfq-bound motifs in three different types of peaks, including strong Hfq-bound peaks, all Hfq-bound peaks and RNA-Seq peaks. Bar height represents the enrichment of motifs (-log10 *p*-value).

**Fig S7. Model presentation of the formation of Hfq-sRNA-mRNA complex, the coupled cleavage of both sRNA and mRNA by RNase E, and the recycling of sRNAs.**

**Table S1. Mapping of clean reads on the reference genome.**

**Table S2. Mapping of clean reads on the reference genome. Reads were classified as the RNA type of CLIP-Seq samples.**

**Table S3. Hfq-bound and –unbound genes. Table S4. Hfq-bound sRNA table.**

**Table S5. Peak summary and classification by region.**

**Table S6. Homology analysis of sRNAs from different studies.**

**Table S7. sRNA table identified from the HFQ_Flag strain. The table merged the CLIP-Seq and RNA-Seq peaks as a union set.**

