## Supplementary figures and images for "Hfq Globally Binds and Destabilizes the bound sRNAs and mRNAs in *Yersinia pestis*"

### Supplementary file 1

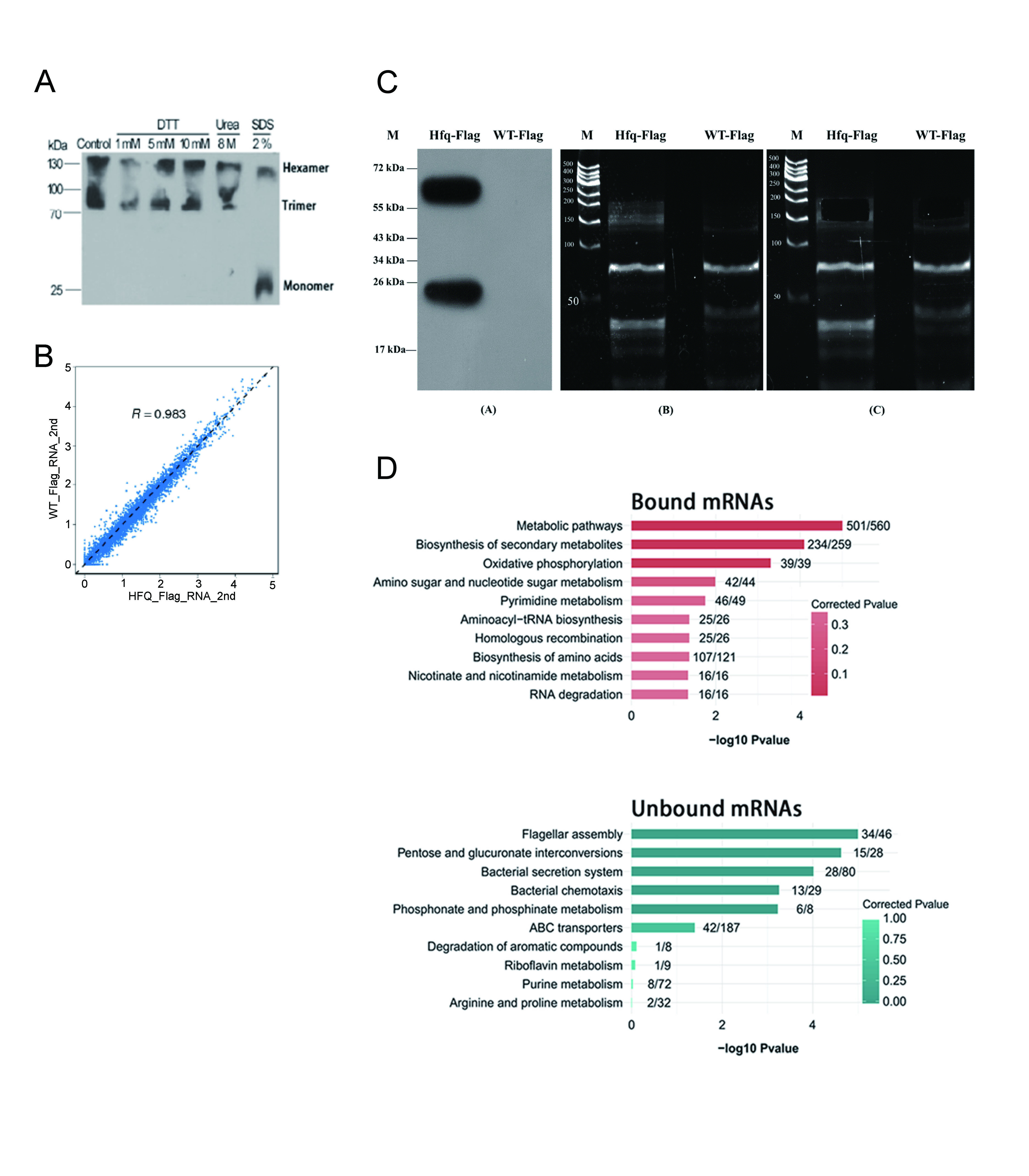

### Supplementary file 3

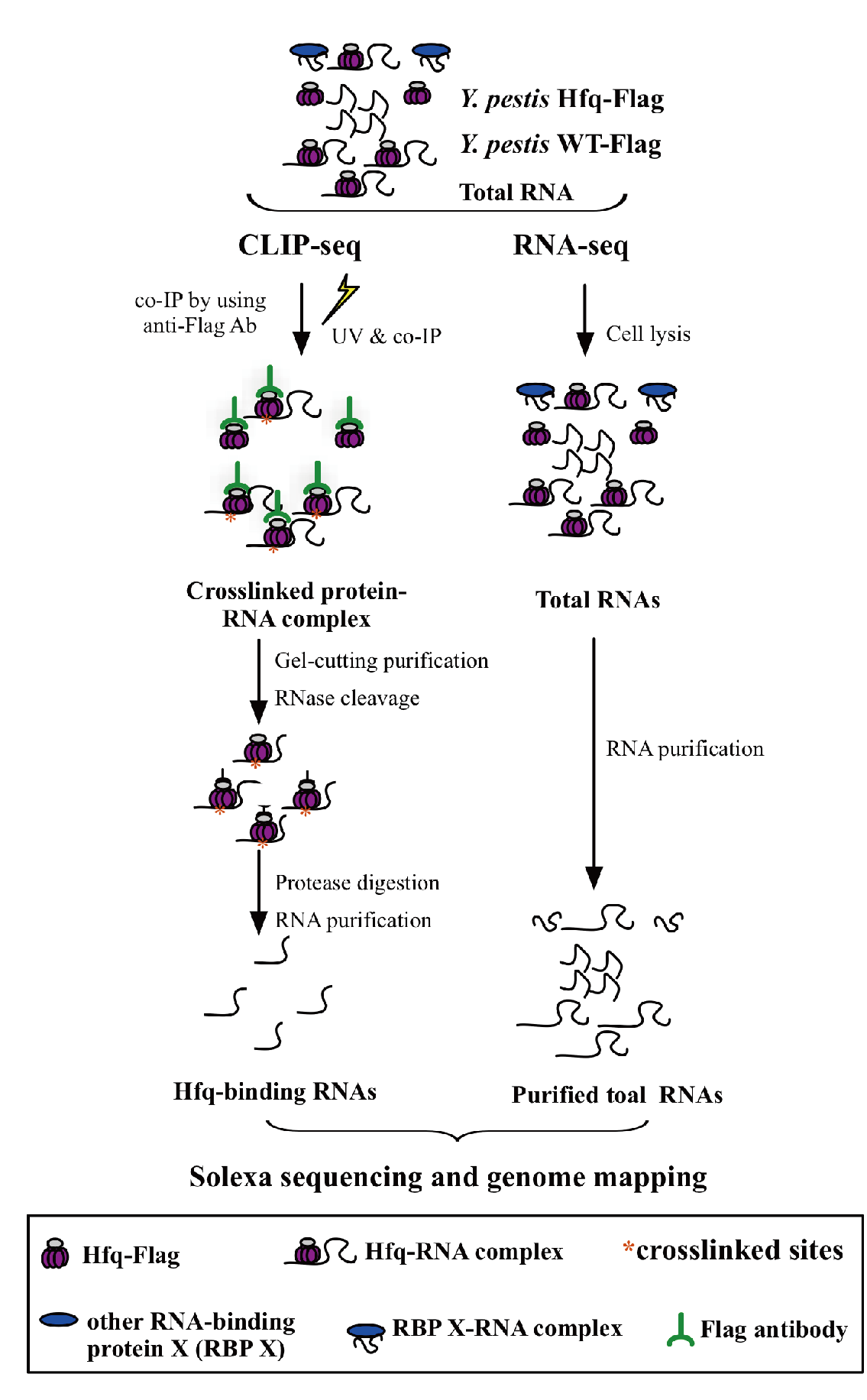

### Supplementary file 5

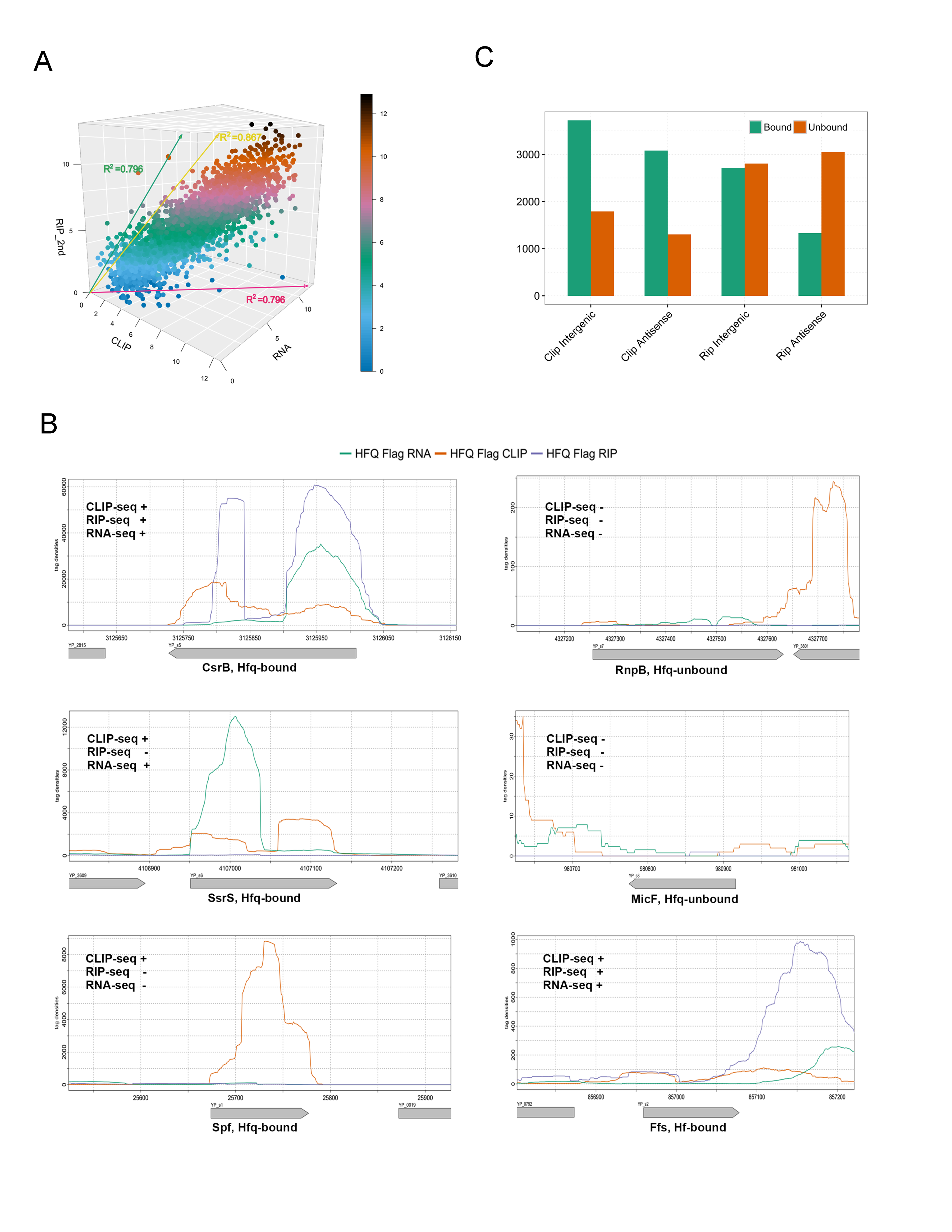

### Supplementary file 7

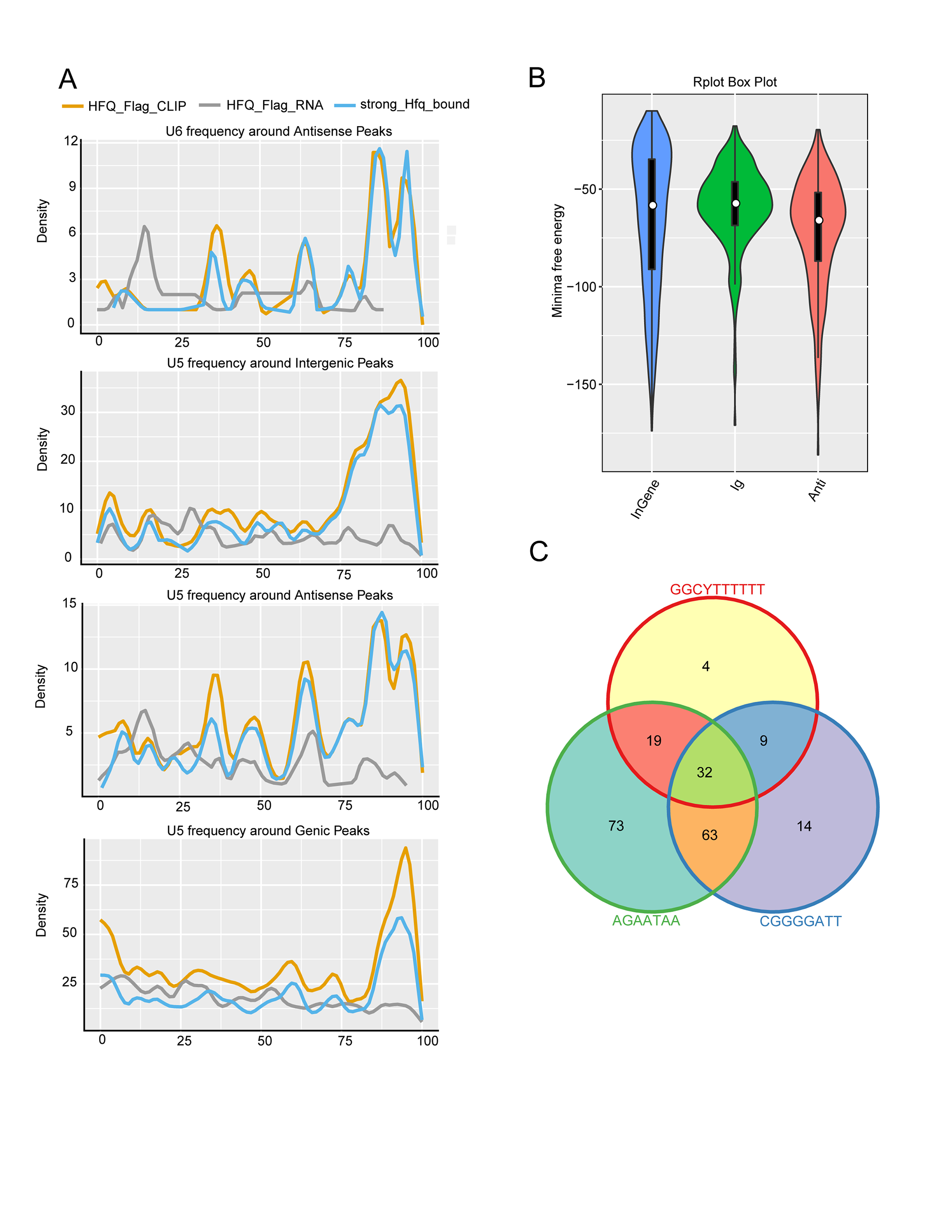

### Supplementary file 9

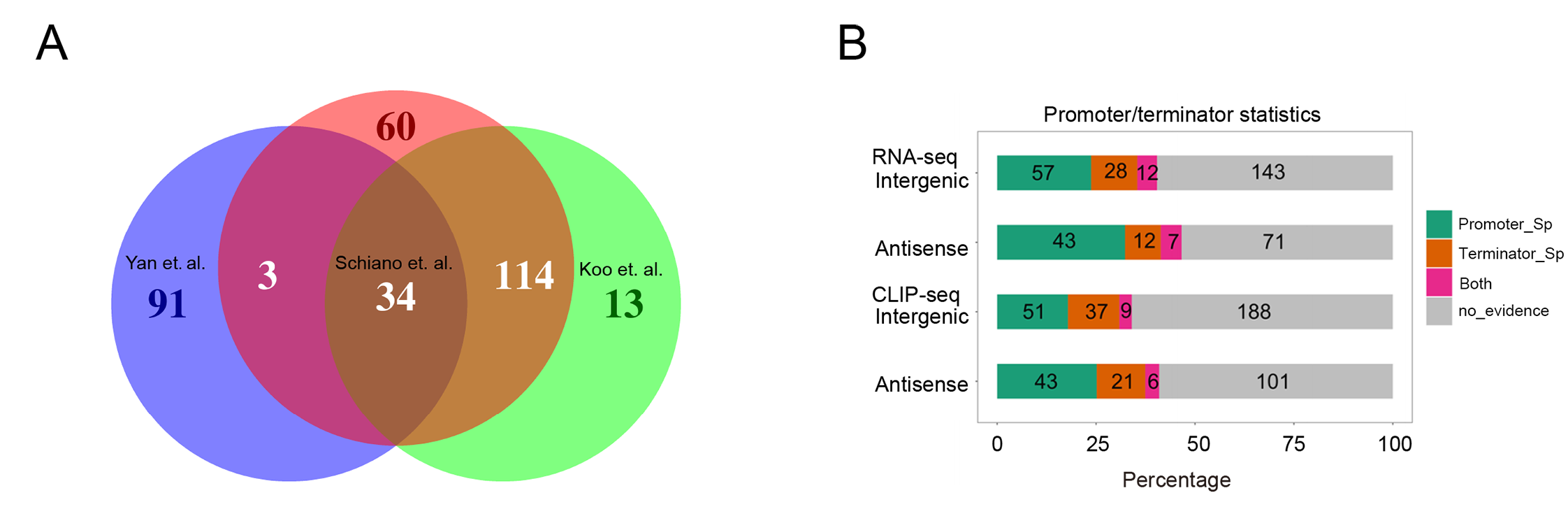

### Supplementary file 11

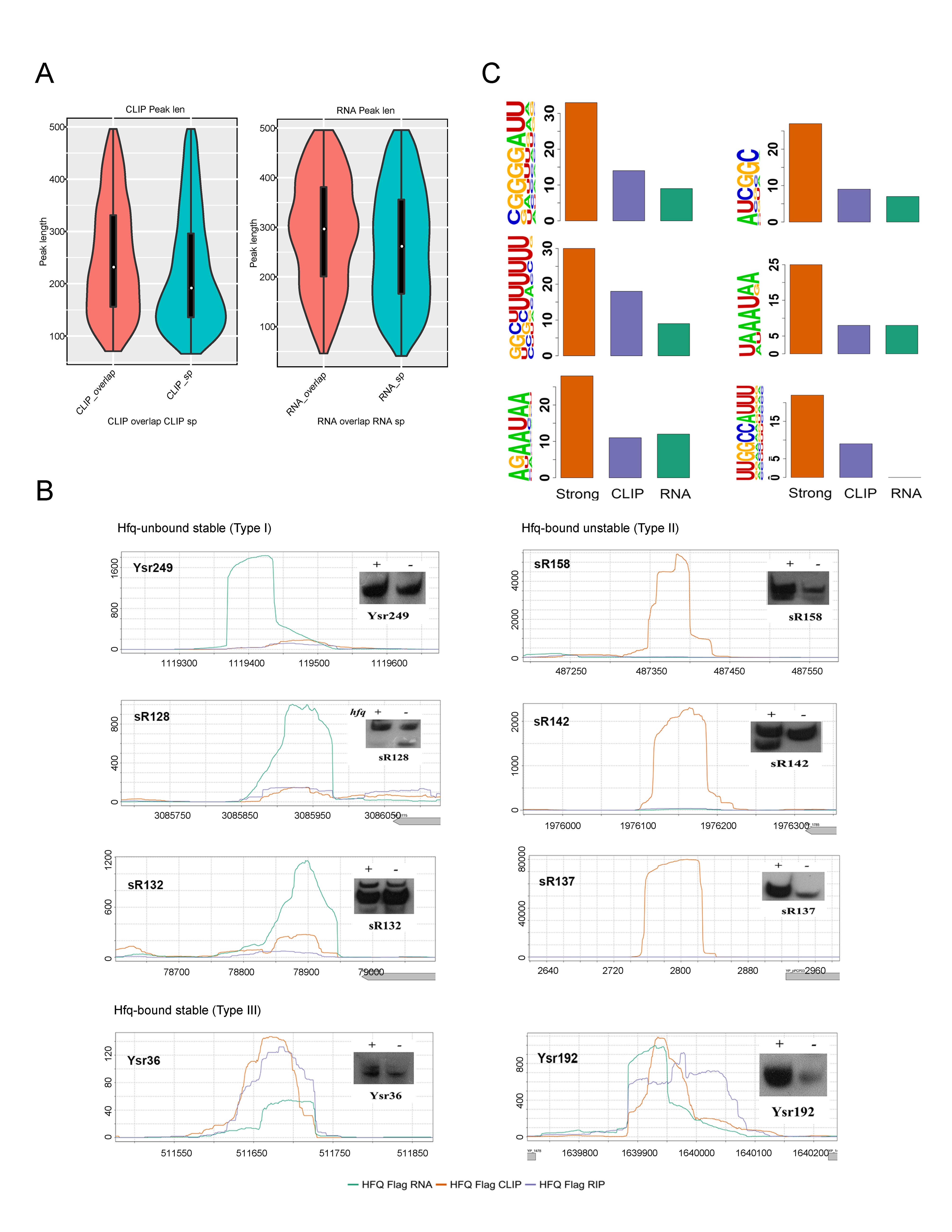

### Supplementary file 13

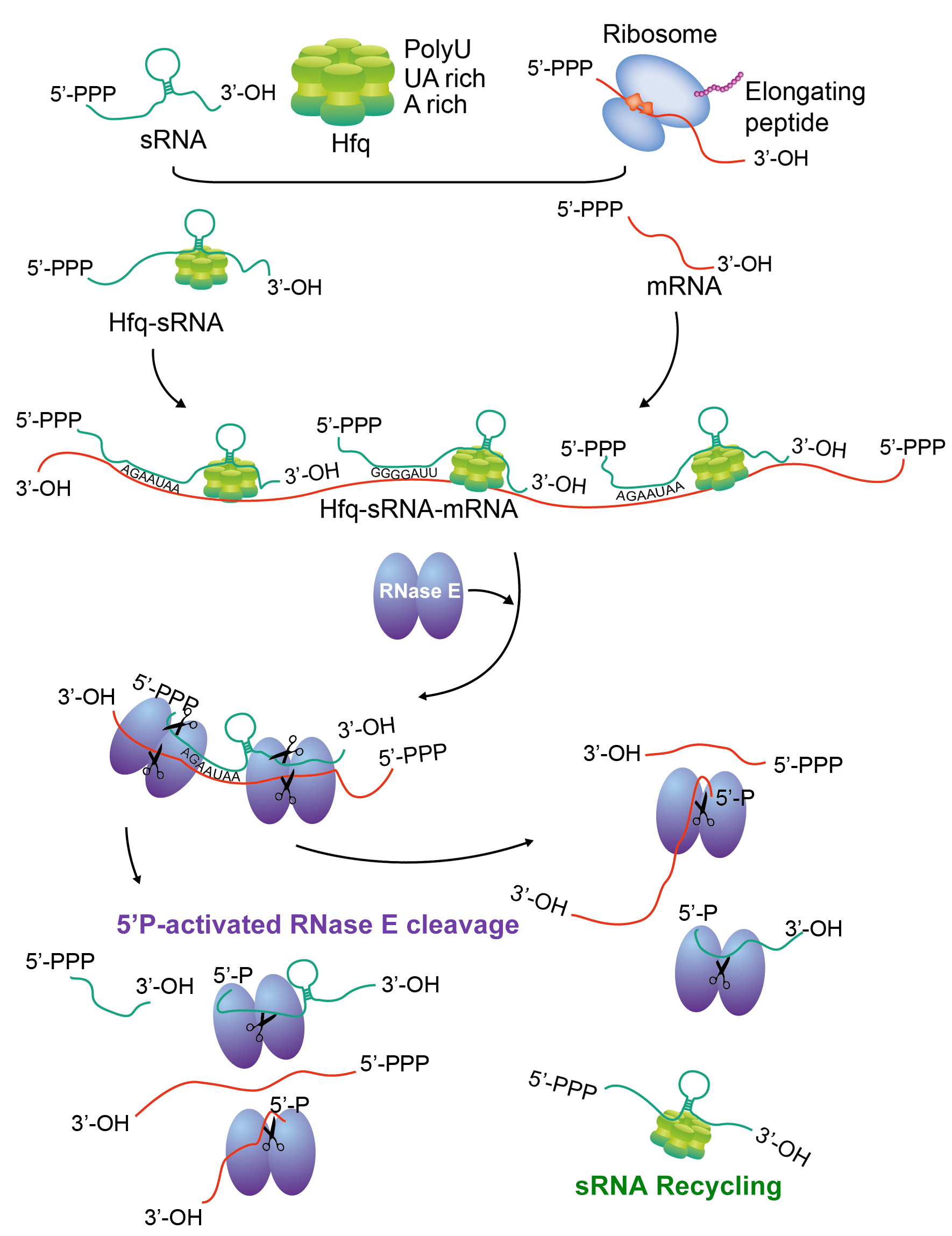
