## Supplementary material for "Hfq Globally Binds and Destabilizes the bound sRNAs and mRNAs in *Yersinia pestis*"

| **Table S1. Mapping of clean reads on the reference genome.** | | | | | |
| --- | --- | --- | --- | --- | --- |
| **Sample** | **input data** | **total mapped** | **uniq mapped** | **multiple mapped** | **antisense** |
| Hfq_FLAG_CLIP | 25,537,735 | 18,023,808  70.58% | 5,871,361  22.99% | 12,152,447  47.59% | 343,770  5.86% |
| WT-FLAG_CLIP | 9,067,353 | 2,065,780  22.78% | 110,743  1.22% | 1,955,037  21.56% | 27,469  24.80% |
| Hfq_FLAG_RIP_2nd | 26,971,805 | 22,624,137  83.88% | 6,602,442  24.48% | 16,021,695  59.40% | 187,051  2.83% |
| FLAG_RIP_2nd | 1,206,503 | 602,544  49.94% | 15,008  1.24% | 587,536  48.70% | 1,972  13.14% |
| WT_FLAG_RIP_1st | 9,186,327 | 6,145,288  66.90% | 798,789  8.70% | 5,346,499  58.20% | 161,633  20.23% |
| WT-FLAG_RIP_1st | 2,291,269 | 1,420,373  61.99% | 56,205  2.45% | 1,364,168  59.54% | 48,425  86.16% |
| Hfq_FLAG_RNA | 7,664,243 | 7,041,344  91.87% | 6,681,618  87.18% | 359,726  4.69% | 291,135  4.36% |
| WT-FLAG_RNAd | 9,631,134 | 8,542,765  88.70% | 7,969,272  82.74% | 573,493  5.95% | 321,528  4.03% |
| WT_RNA | 6,306,946 | 5,680,049  90.06% | 5,220,651  82.78% | 459,398  7.28% | 191,022  3.66% |
