## Supplementary material for "Hfq Globally Binds and Destabilizes the bound sRNAs and mRNAs in *Yersinia pestis*"

| **Table S2. Mapping of clean reads on the reference genome. Reads were classified as the RNA type of CLIP-Seq samples.** | | | | | | | |
| --- | --- | --- | --- | --- | --- | --- | --- |
| **Sample** | **input data** | **total mapped** | **mRNA** | **sRNA** | **rRNA** | **tRNA** | **Intergenic RNA** |
| Hfq_FLAG_CLIP | 25,537,735 | 18,023,808 | 3,655,093 | 54,239 | 11,697,891 | 25,694 | 2,590,891 |
| Percentage | 100% | 70.58% | 20.28% | 0.30% | 64.90% | 0.10% | 14.37% |
| WT-FLAG_CLIP | 9,067,353 | 2,065,780 | 51,283 | 649 | 1,777,134 | 215,826 | 20,888 |
| Percentage | 100% | 22.78% | 2.48% | 0.03% | 86.03% | 2.38% | 1.01% |
