## Supplementary material for "Hfq Globally Binds and Destabilizes the bound sRNAs and mRNAs in *Yersinia pestis*"

| **Table S4. Hfq-bound sRNA table, Related to Figure 1.** | | | | | | | | | |
| --- | --- | --- | --- | --- | --- | --- | --- | --- | --- |
| **#Gene** | **Chromsome** | **WT-FLAG** | | **HFQ_FLAG** | | **Fold enriched** | **Gene name** | **This study** | **Previous Studies*** |
| **sRNAs** |  | **Reads** | **Ratio** | **Reads** | **Ratio** |  |  |  |  |
| YP_s1 | NC_005810.1 | 163 | 0.00069 | 9472 | 0.00560 | 8.15 | *spf* | Hfq-bound | Bound in E. coli and Salmonella |
| YP_s2 | NC_005810.1 | 1 | 0.00000 | 92 | 0.00005 | 12.90** | *Ffs*** | Hfq-unbound | Unbound in E. coli |
| YP_s3 | NC_005810.1 | 0 |  | 0 |  | NA | *micF* | No expression | Bound in E. coli and Salmonella |
| YP_s4 | NC_005810.1 | 3 | 0.00001 | 2 | 0.00000 | 0.09 | *ssrA* | Hfq-unbound | Unbound in E. coli |
| YP_s5 | NC_005810.1 | 222 | 0.00094 | 37637 | 0.02224 | 23.78 | *csrB* | Hfq-bound | Unbound in Salmonella |
| YP_s6 | NC_005810.1 | 253 | 0.00107 | 5631 | 0.00333 | 3.12 | *ssrS* | Hfq-bound | Unbound in E. coli and Salmonella |
| YP_s7 | NC_005810.1 | 2 | 0.00001 | 25 | 0.00001 | 1.75 | *rnpB* | Hfq-unbound | Unbound in E. coli |
| pCD06.1 | NC_005813.1 | 5 | 0.00002 | 1380 | 0.00082 | 38.71 | *pCD06.1* | Hfq-bound | NA |
| Control |  |  |  |  |  |  |  |  |  |
| YP_r2 | NC_005810.1 | 237382 | 1.00000 | 1692371 | 1.00000 | 1.00 | YP_r2 (23S rRNA) | |  |

*Reference for Hfq-bound sRNA in *E. coli*: Zhang et al., 2003, Molecular Microbiology 50:1111-1124

*Reference for Hfq-bound sRNA in *Salmonella*: Sittka et al., 2008, Plos Genetics e1000163

**Please be noted the *Ffs* sRNA annotation shown here was according to the previously published genome annotation, which was shown to be wrong in the strand direction. The correct *Ffs* sRNA on the opposite direction was among the sRNA peaks identified from RNA-seq data (Table S7) and shown to be Hfq-unbound (Figure 6F).
