## Supplementary material for "Hfq Globally Binds and Destabilizes the bound sRNAs and mRNAs in *Yersinia pestis*"

| **Table S5-Peak summary and classification by region.** | | | | | |
| --- | --- | --- | --- | --- | --- |
| **Sample** | **Total** | **InGene** | **Intergenic** | **Antisense** | **Asso gene** |
| HFQ_Flag_CLIP | 2511 | 2055 | 285 | 171 | 1619 |
| WT_Flag_CLIP | 46 | 27 | 16 | 3 | 27 |
| HFQ_Flag_RNA | 1518 | 1268 | 164 | 86 | 1095 |
| WT_Flag_RNA | 1731 | 1416 | 206 | 109 | 1154 |
| HFQ_RNA | 1447 | 1209 | 159 | 79 | 987 |
