## Supplementary material for "Hfq Globally Binds and Destabilizes the bound sRNAs and mRNAs in *Yersinia pestis*"

| **Table S6. Homology analysis of sRNAs from different studies.** | | | | | | |
| --- | --- | --- | --- | --- | --- | --- |
| **Previous study** | **This study** | **Previous-study Aligned** | | | **This-study Aligned** | |
| Yan et al. 2013 (117) | RNA-Seq (373) | 50 | 42.74% | 65 | | 17.42% |
| Yan et al. 2013 (117) | CLIP-Seq (456) | 55 | 47.01% | 62 | | 13.60% |
| Schiano et al., 2014 (211) | RNA-Seq (373) | 83 | 39.34% | 78 | | 20.91% |
| Schiano et al., 2014 (211) | CLIP-Seq (456) | 97 | 45.97% | 86 | | 18.86% |
| Koo et al., 2011 (148) | RNA-Seq (373) | 63 | 41.89% | 62 | | 16.62% |
| Koo et al., 2011 (148) | CLIP-Seq (456) | 73 | 49.32% | 70 | | 15.35% |
